# Viral evolution prediction identifies broadly neutralizing antibodies against existing and prospective SARS-CoV-2 variants

**DOI:** 10.1101/2024.04.16.589454

**Authors:** Fanchong Jian, Anna Z. Wec, Leilei Feng, Yuanling Yu, Lei Wang, Peng Wang, Lingling Yu, Jing Wang, Jacob Hou, Daniela Montes Berrueta, Diana Lee, Tessa Speidel, LingZhi Ma, Thu Kim, Ayijiang Yisimayi, Weiliang Song, Jing Wang, Lu Liu, Sijie Yang, Xiao Niu, Tianhe Xiao, Ran An, Yao Wang, Fei Shao, Youchun Wang, Simone Pecetta, Xiangxi Wang, Laura M. Walker, Yunlong Cao

## Abstract

Monoclonal antibodies (mAbs) targeting the SARS-CoV-2 receptor-binding domain (RBD) are used to treat and prevent COVID-19. However, the rapid evolution of SARS-CoV-2 drives continuous escape from therapeutic mAbs. Therefore, the ability to identify broadly neutralizing antibodies (bnAbs) against future variants is needed. Here, we use deep mutational scanning (DMS) to predict viral RBD evolution and to select for mAbs neutralizing both existing and prospective variants. A retrospective analysis of 1,103 SARS-CoV-2 wildtype-elicited mAbs shows that this method can increase the probability of identifying effective bnAbs against the XBB.1.5 strain from 1% to 40% in an early pandemic setup. Among these bnAbs, BD55-1205 exhibited potent activity against all tested variants. Cryo-EM structural analyses revealed the receptor mimicry of BD55-1205, explaining its broad reactivity. Delivery of mRNA-LNPs encoding BD55-1205-IgG in mice resulted in ~5,000 serum NT_50_ against XBB.1.5, HK.3.1, and JN.1 variants. Combining bnAb identification using viral evolution prediction with the versatility of mRNA delivery technology can enable rapid development of next-generation antibody-based countermeasures against SARS-CoV-2 and potentially other pathogens with pandemic potential.

## Introduction

Severe acute respiratory syndrome coronavirus 2 (SARS-CoV-2) continues to rapidly evolve to evade immunity induced by natural infection and vaccination, resulting in highly evasive variant lineages like XBB.1.5 and JN.1 ^1,2^. These variants are continuously accumulating mutations at key RBD antigenic sites, such as L455, F456, and A475, which may significantly alter their antigenicity and further escape neutralizing antibodies (NAbs) elicited by repeated vaccination and infection ^3–7^.

Monoclonal NAbs targeting the SARS-CoV-2 RBD have shown high efficacy in the treatment and prevention of COVID-19, especially in high-risk individuals who do not mount robust immune responses to vaccination ^8–12^. However, all previously approved anti-SARS-CoV-2 mAbs and cocktail therapeutics, discovered before the emergence of variants of concern (VOCs), have lost effectiveness against contemporary variants ^13–16^. Since the emergence and rapid evolution of Omicron in 2021, numerous antibodies have been reported to be “broadly neutralizing” or “variant proof” based on their ability to neutralize historical SARS-CoV-2 variants, targeting either RBD, N-terminal domain (NTD), or subdomain 1 (SD1) of the virus Spike glycoprotein ^17–24^. Unfortunately, the vast majority of them rapidly lost activity against newly emerged variants, raising questions about the criteria used to designate antibodies as broadly neutralizing and undermining confidence in the development of next-generation antibodies against SARS-CoV-2 ^14,25,26^. Therefore, the development of a practical strategy to identify bnAbs with neutralizing activity against both existing and prospective variants would greatly enhance the feasibility of developing future antibody-based countermeasures against SARS-CoV-2 that can outpace viral evolution, and would have been invaluable at the early stages of a pandemic.

Previously, we employed high-throughput deep mutational scanning (DMS) on extensive panels of mAbs to characterize the evolutionary pressures on the SARS-CoV-2 RBD to predict future mutation hotspots ^25,27^. Here, we demonstrate that DMS-based mutation prediction can significantly enhance the probability of identifying bnAbs that potently neutralize both existing and future variants. Using this method, we identified a human IGHV3-66-derived Class 1 bnAb elicited by SARS-CoV-2 ancestral strain infection, designated as BD55-1205, which demonstrates extraordinary neutralization breadth against all existing variants as well as prospective variants with mutations within its targeting epitope. Delivery of mRNA-encoded BD55-1205 IgG in mice resulted in high serum neutralizing titers against evasive variants, providing evidence that mRNA technology could be leveraged for rapid deployment of anti-SARS-CoV-2 bnAbs. This platform may enable the rapid development of next-generation antibody-based countermeasures against SARS-CoV-2, and potentially future pandemic pathogens.

## Results

### Retrospective assessment of SARS-CoV-2 NAbs

To date, anti-SARS-CoV-2 NAbs potent against currently circulating variants have been selected for clinical development. However, such NAbs have been repeatedly escaped by Omicron variants, suggesting that activity against current variants does not translate into breadth against future variants^13,16,25^. To investigate the relationship between neutralization potency and breadth against SARS-CoV-2 variants, we studied a panel of 7,018 mAbs isolated from 7 previously described cohorts, which include individuals infected or vaccinated by ancestral SARS-CoV-2 (hereafter denoted as “WT”), individuals who experienced SARS-CoV-1 infection in 2003/2004 and received 3-dose CoronaVac in 2021 (SARS+WT), convalescents who experienced BA.1, BA.2, or BA.5/BF.7 breakthrough infection (BTI) after 3-dose CoronaVac (BA.1 BTI, BA.2 BTI, BA.5/BF.7 BTI, respectively), and convalescents who experienced BA.1 or BA.2 BTI and were reinfected by BA.5/BF.7 (BA.1 or BA.2 BTI+BA.5/BF.7) ^13,14,25,27,28^. Out of the 7,018 mAbs, we identified 1,637 potent autologous NAbs, defined as IC_50_ < 0.05 μg/mL against the corresponding last-exposure variant (Fig. 1a and Table S1).

**Figure 1.**
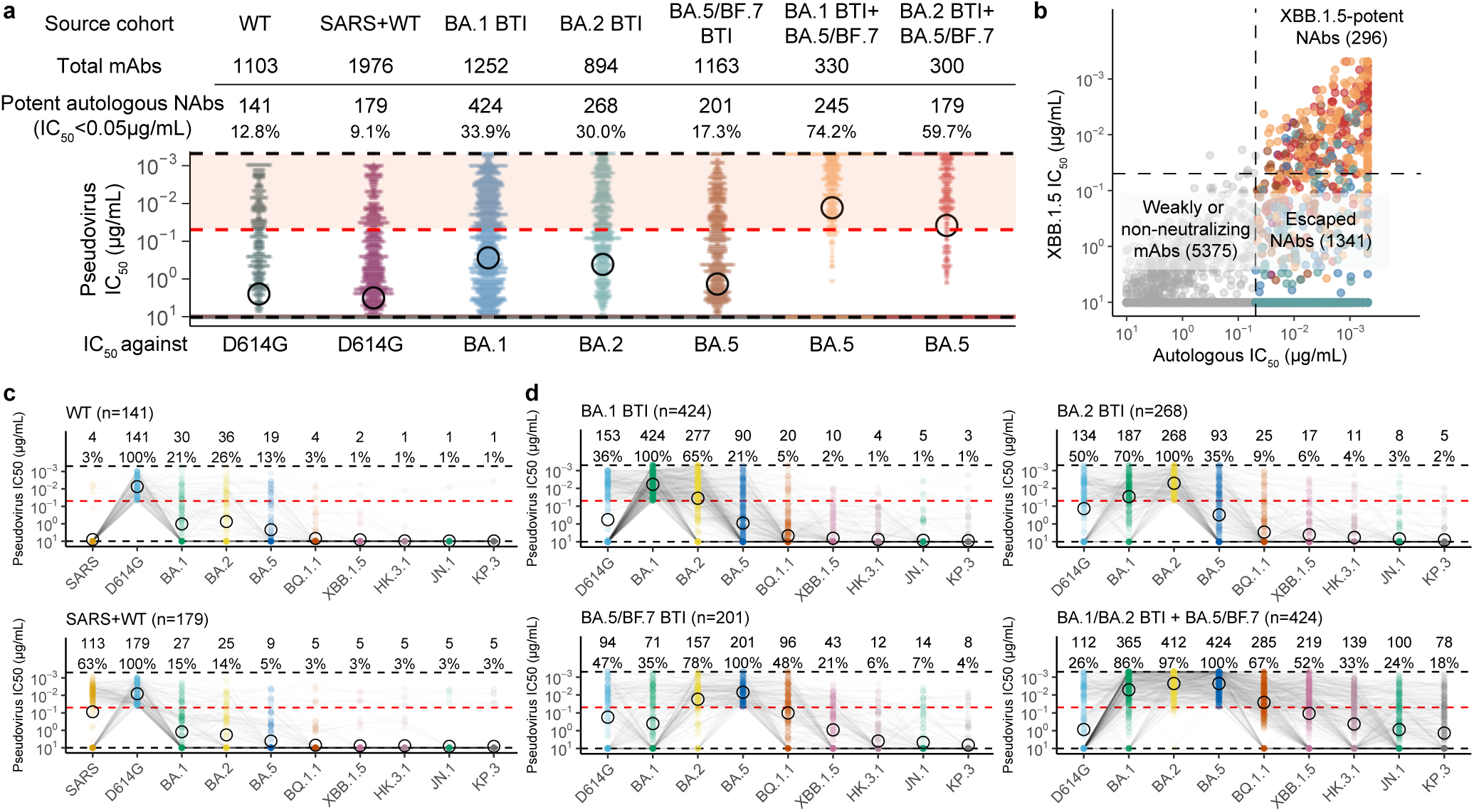
Neutralization activities of RBD-targeting mAbs against SARS-CoV-2 variants. **a**, Neutralization of the 7,018 mAbs from individuals with 7 different immune histories against the corresponding last-exposure variant (autologous neutralization activity). Numbers and proportions of potent autologous NAbs (IC_50_ < 0.05 μg/mL) are annotated above each group of points. The black circles indicate the geometric mean values of each group. **b**, Relationship between the autologous neutralization activities and XBB.1.5-neutralizing activities of the isolated mAbs. **c-d**, Neutralization activities of potent autologous NAbs against variants. Numbers and proportions of potent NAbs against each variant (IC_50_ < 0.05 μg/mL) are annotated.

To systematically investigate the breadth of NAbs isolated from various cohorts, we tested the neutralization activities of the 1,637 potent autologous NAbs against eight major SARS-CoV-2 variants, including B.1 (D614G), Omicron BA.1, BA.2, BA.5, BQ.1.1, XBB.1.5, HK.3, and JN.1 (Fig. 1b-d). Among the 1,637 potent NAbs from 6 cohorts, only 296, 133, and 100 mAbs demonstrated potent neutralization against XBB.1.5, JN.1, and KP.3, respectively (Fig. 1b and Extended Data Fig. 1a). Although there was an association between autologous and XBB.1.5, JN.1 or KP.3 neutralization activity, NAbs with the strongest autologous neutralization activities generally lost neutralization breadth against “future” variants (at that time); this phenomenon was particularly striking for NAbs obtained at the early stages of the pandemic from individuals infected or vaccinated with the ancestral strain (Fig. 1b and Extended Data Fig. 1b). Among the 141 WT-elicited potent NAbs, only two (1%) remained potent against XBB.1.5, and only one potently neutralized all tested variants including JN.1 (Fig. 1c). BA.1 and BA.2-elicited antibodies exhibited slightly better tolerance to subsequent Omicron subvariants, but nevertheless, only 10/424 (2%) and 17/268 (7%) showed activity against XBB.1.5, respectively (Fig. 1d). Although a higher proportion of mAbs from the BA.5/BF.7 BTI or reinfection cohorts neutralized BQ.1.1 and XBB.1.5, only 7% and 27% potently neutralized JN.1, respectively (Fig. 1d). The vast majority (85%) of SARS-CoV-1/SARS-CoV-2 cross-reactive mAbs isolated from the SARS+WT cohort showed abrogated activity against Omicron variants (Fig. 1c) ^13,28^. Only five out of the 179 autologous potent NAbs from SARS+WT potently neutralized all tested variants, and all of them belong to epitope group F3, similar to SA55 ^28^. Therefore, sarbecovirus-based identification of bnAbs does not guarantee neutralization breadth against emerging SARS-CoV-2 variants, and neutralization potency against circulating variants at the time of isolation is not a sufficient metric to define neutralization breadth. Thus, a generalizable strategy to accurately identify bnAbs that retain activity against future SARS-CoV-2 variants is of paramount importance for the development of next-generation NAb-based therapeutics.

### The rational strategy to select bnAbs

Previously, we demonstrated that the integration of DMS profiles accurately predicted positively selected mutations during viral evolution under humoral immune pressure ^25,27^. We hypothesized that screening mAbs against pseudoviruses encoding predicted mutations could identify bnAbs resilient to future variants efficiently.

To validate this strategy, we retrospectively studied a collection of the mAbs elicited by SARS-CoV-2 WT exposure, which could be obstained early in the pandemic ^29,30^. We integrated antibody DMS escape profiles, codon preferences, human ACE2 (hACE2) binding and RBD expression impacts to map hotspots on RBD, including R346, K378, K417, K444-G446, N450, L452, E484, F486, F490, and Q493 (Fig. 2a) ^25,27,31^. We constructed 17 mutant pseudoviruses harboring single amino acid substitutions at these positions based on B.1 (D614G) strain (Extended Data Fig. 2a). Because E484K is the most striking mutation from the calculation, we first tested the neutralization activities of 141 potent NAbs from the WT cohort against B.1+E484K. For mAbs remaining effective, the neutralization activities against the other 16 single-substitution mutants were tested. These mutants can help enrich for Omicron-effective NAbs, increasing the frequency of BA.1/BA.2/BA.5-effective NAbs from 14/141 (9.9%) to 12/46 (26%) (Extended Data Fig. 2b). To enhance the efficiency to escape antibodies, we further designed five mutants with combination of these mutations, named B.1-S1 to S5 (Fig. 2b). The combinations were designed to maintain hACE2 binding and avoid multiple mutations in the same epitope (Supplementary Information). As expected, these mutants, especially B.1-S3 (R346T+K417T+K444N+L452R+E484K+F486S), substantially escaped the majority of potent NAbs elicited by WT exposure (Fig. 2c and Extended Data Fig. 2a). Passing the filter of all five design mutants (S1-S5 refers to the weakest neutralization against the 5 designed mutants) is shown to be an efficient indicator of broad neutralization against real-world Omicron variants (Fig. 2d-e). NAbs potently neutralizing all of B.1-S1 to S5 significantly enriches true bnAbs capable of neutralizing Omicron subvariants (Fig. 2e).

**Figure 2.**
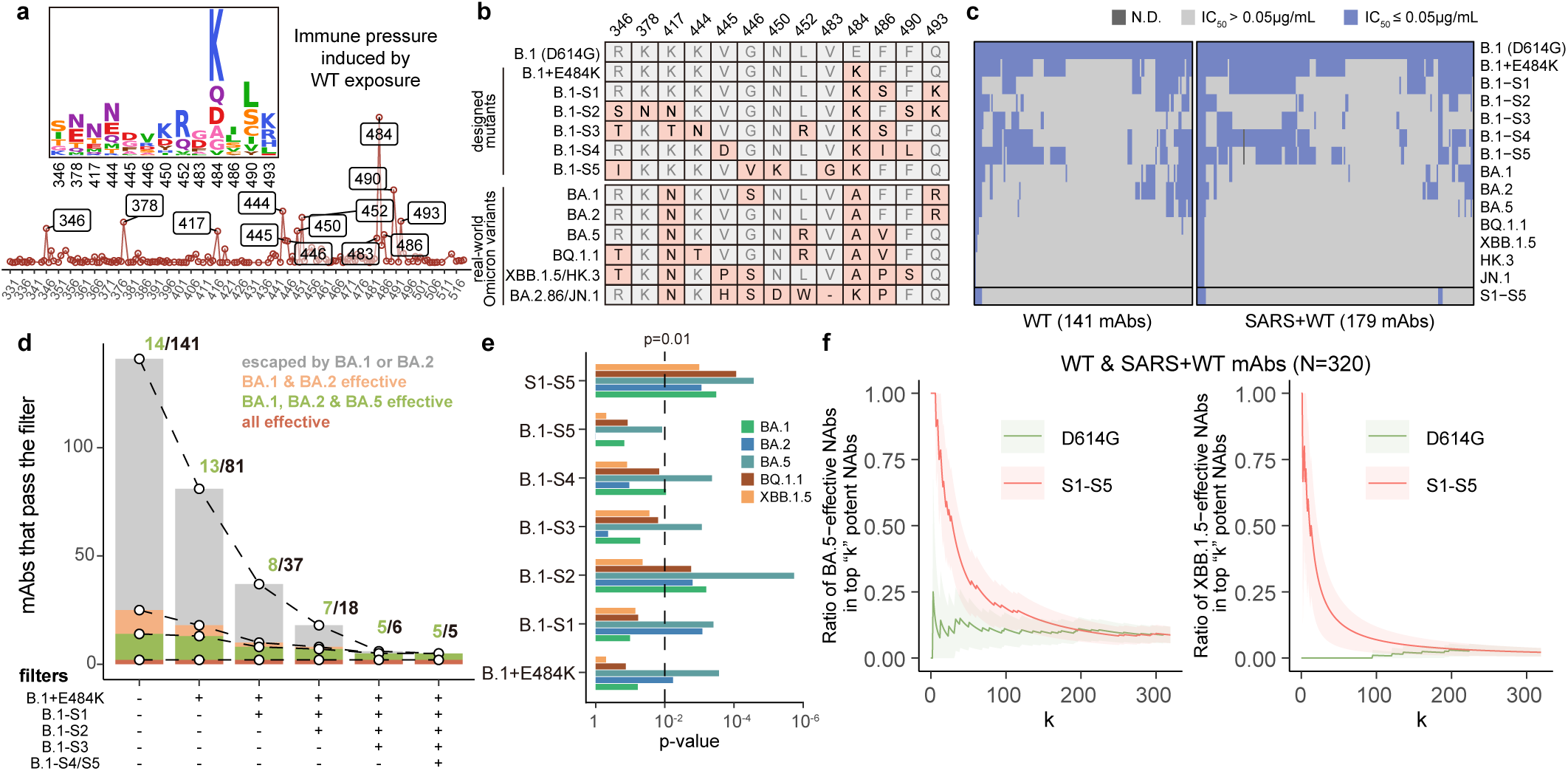
Designed mutants based on mutation prediction define bnAbs. **a**, Average escape profiles from DMS of mAbs (weighted by neutralization activities of each mAb against SARS-CoV-2 B.1 and the impact of each RBD mutation on ACE2 binding and RBD expression). Y-axis corresponds to weighted average escape scores across all mAbs on each RBD residue (relative values). **b**, Mutations harbored by the designed SARS-CoV-2 B.1-based mutants and real-world variants on the key sites indicated by DMS-based prediction. **c**, Neutralization capability of the mAbs from early cohorts (SARS+WT and WT) against the designed mutants and real-world Omicron variants. “S1-S5” indicates the highest IC_50_ against the five designed mutants. **d**, Number of NAbs from WT vaccinees or convalescents that pass the filter of designed mutants. Ratio of BA.1, BA.2, and BA.5-potent NAbs among the passed NAbs are annotated above the bar of each combination of filters. **e**, Significance for the enrichment of BA.1, BA.2, BA.5, BQ.1.1, or XBB.1.5-potent NAbs within NAbs that are from WT vaccinees or convalescents and pass each filter of designed mutants (two-sided hypergeometric test). **f**, Ratio of BA.5 or XBB.1.5-potent NAbs within the NAbs with “top k” neutralization activities against D614G or S1-S5. The error bars indicate 95% confidence interval under normal distribution.

Only 5 out of 141 NAbs from WT remained potent (IC_50_ < 0.05 μg/mL) against all of the rationally designed evasive mutants (Fig. 2d). Notably, although the mutations of our predicted mutant sequences only partially overlap with real-world Omicron variants, all 5 NAbs with activity against all rationally designed mutants effectively neutralized Omicron BA.1, BA.2, and BA.5. Importantly, the five selected bnAbs include the only two XBB.1.5-effective antibodies among the whole collection of 141 candidates, which corresponds to an increase in the probability of identifying “true” bnAbs from ~1% to 40%. Overall, our model correctly predicted the key escape mutations incorporated during SARS-CoV-2 antigenic drift and allowed for the identification of a small subset of WT-elicited bnAbs with activity against future variants.

Additionally, we analyzed the panel of 320 NAbs from WT (n=141) and SARS+WT (n=179) cohorts, all of which could have been obtained early in the pandemic. Similarly, the neutralization activities of the SARS+WT-elicited NAbs against the designed mutants predicted BA.5/XBB.1.5 activity (Extended Data Fig. 2b-c). In contrast, most potent B.1 NAbs were escaped by XBB.1.5/JN.1, underscoring the value of designed mutants (Fig. 2f and Extended Data Fig. 2d-e). The results validate our viral evolution prediction platform for the identification of rare, resilient bnAbs from a large collection of antibodies isolated from convalescent or vaccinated individuals at the early stage of a pandemic.

### BD55-1205 exhibits high barrier to escape

BD55-1205, a Class 1 public antibody that utilizes the IGHV3-53/3-66 germline ^29,32,33^, was the only JN.1-effective bnAb from the WT cohort (Table 1). It exhibits the strongest “S1-S5” neutralization (refers to the weakest in five designs), which may reflect its better tolerance to mutations compared to other competitors in epitope group A1. Given BD55-1205 exhibited broad neutralization against all variants tested, we assessed whether it also demonstrated high barrier to escape under stringent *in vitro* conditions. Using XBB.1.5 S-pseudotyped replication-competent recombinant vesicular stomatitis virus (rVSV), we screened for antibody escape mutations via serial passage in Vero cells (Fig. 3a and Extended Data Fig. 3a) ^34,35^. This assay simulated the viral evolution under the pressure of a specific mAb. For comparison, a bnAb SA55 and a non-competing pair of XBB.1.5-effective bnAbs (BD57-1520+BD57-2225) were included (Extended Data Fig. 3b-d) ^27^. Surprisingly, BD55-1205 displayed similar resistance to viral escape as the bnAb cocktail, retaining neutralization until passage 6 (Fig. 3b and Extended Data Fig. 4d-e). In contrast, individual SA55, BD57-2225 or BD57-1520 was escaped after two or three passages (Fig. 3b and Extended Data Fig. 4a-c). Substitutions L455P, Q493R/K, N417K, A435T, and D420Y were enriched in the BD55-1205-selected virus, consistent with critical residues interacting with Class 1 antibodies, as indicated by DMS based on BA.5 or XBB.1.5 RBD (Fig. 3c and Extended Data Fig. 3c-d). To validate the selected mutations, we constructed XBB.1.5+L455P+Q493R, XBB.1.5+N417K+A435T, and XBB.1.5+D420Y+Q493K mutant pseudoviruses. The activities of BD55-1205 against them are reduced but not completely escaped (Fig. 3d). Neutralization assays using soluble hACE2 indicated lower receptor binding capability of these mutants compared to XBB.1.5 (Fig. 3d and Extended Data Fig. 5a-b), suggesting that the mutations abrogating BD55-1205 activity may result in reduced viral fitness through disruption of receptor binding interactions.

**Figure 3.**
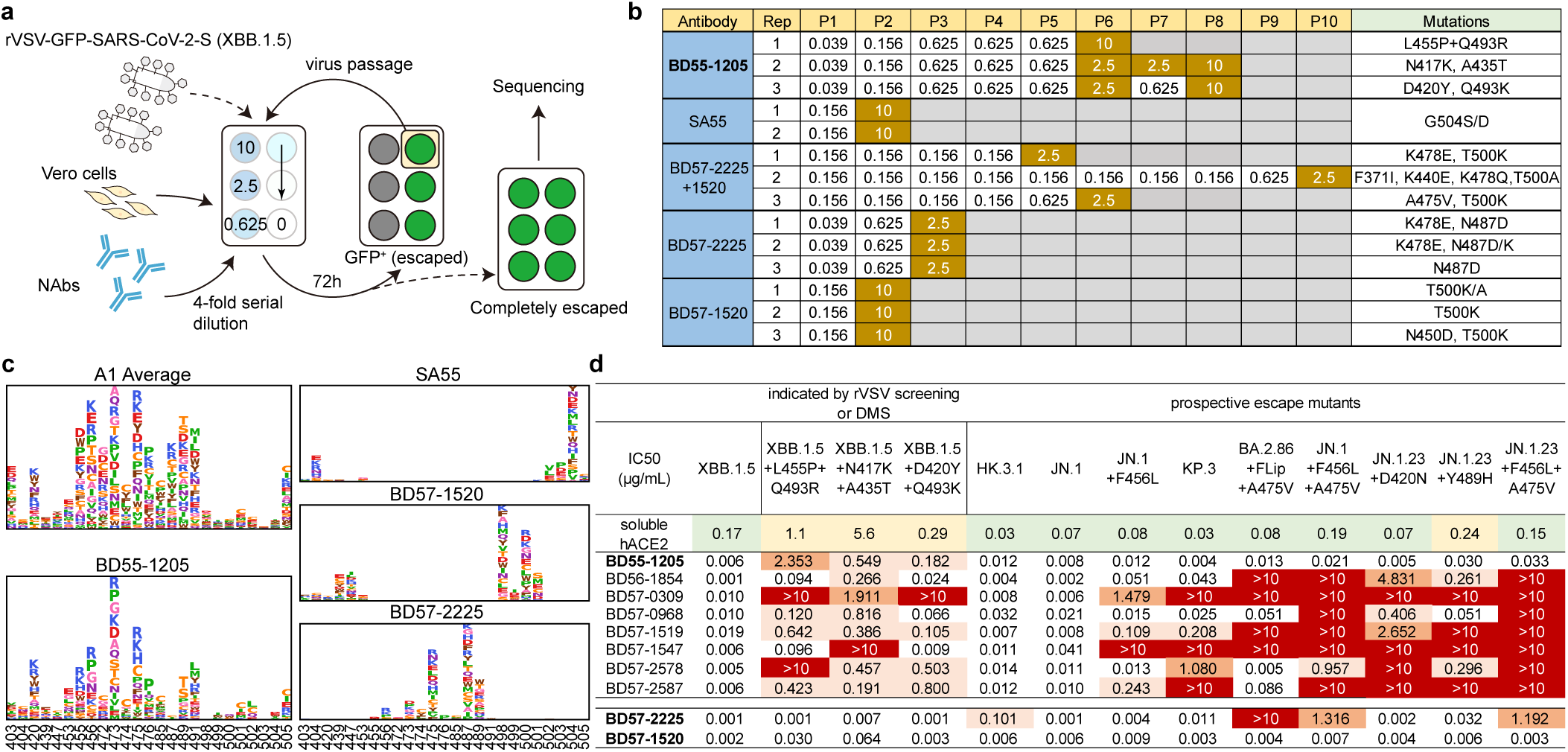
BD55-1205 exhibits extraordinary resistance to viral escape. **a**, Schematic for the rVSV-based escape mutation screening assays. **b**, Results of the escape screening by rVSV passaging under the NAb pressure. Values in the P1-P10 columns of the table indicate the highest concentration of NAb that was escaped by rVSV in the first passage to the tenth passage. The mutations observed in the final passage of rVSV were determined by Sanger sequencing and are annotated in the last column. **c**, DMS escape profiles (based on XBB.1.5 RBD) of the NAbs evaluated in the rVSV assays. The average profile of antibodies in epitope group A1 is also shown for comparison with BD55-1205. **d**, Neutralization of BD55-1205 and other NAbs against designed escape mutants according to rVSV screening and DMS profiles, and real-world emerging and prospective mutants with mutations in the BD55-1205 epitope.

**Table 1.**
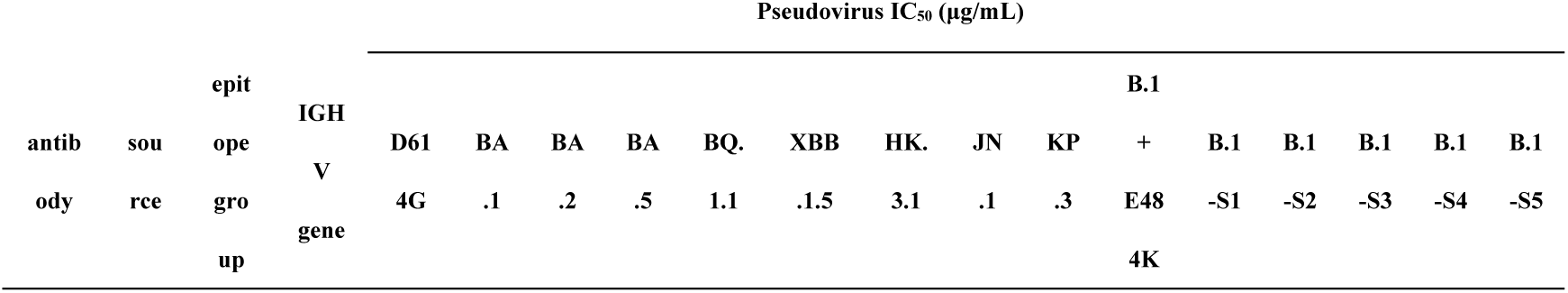
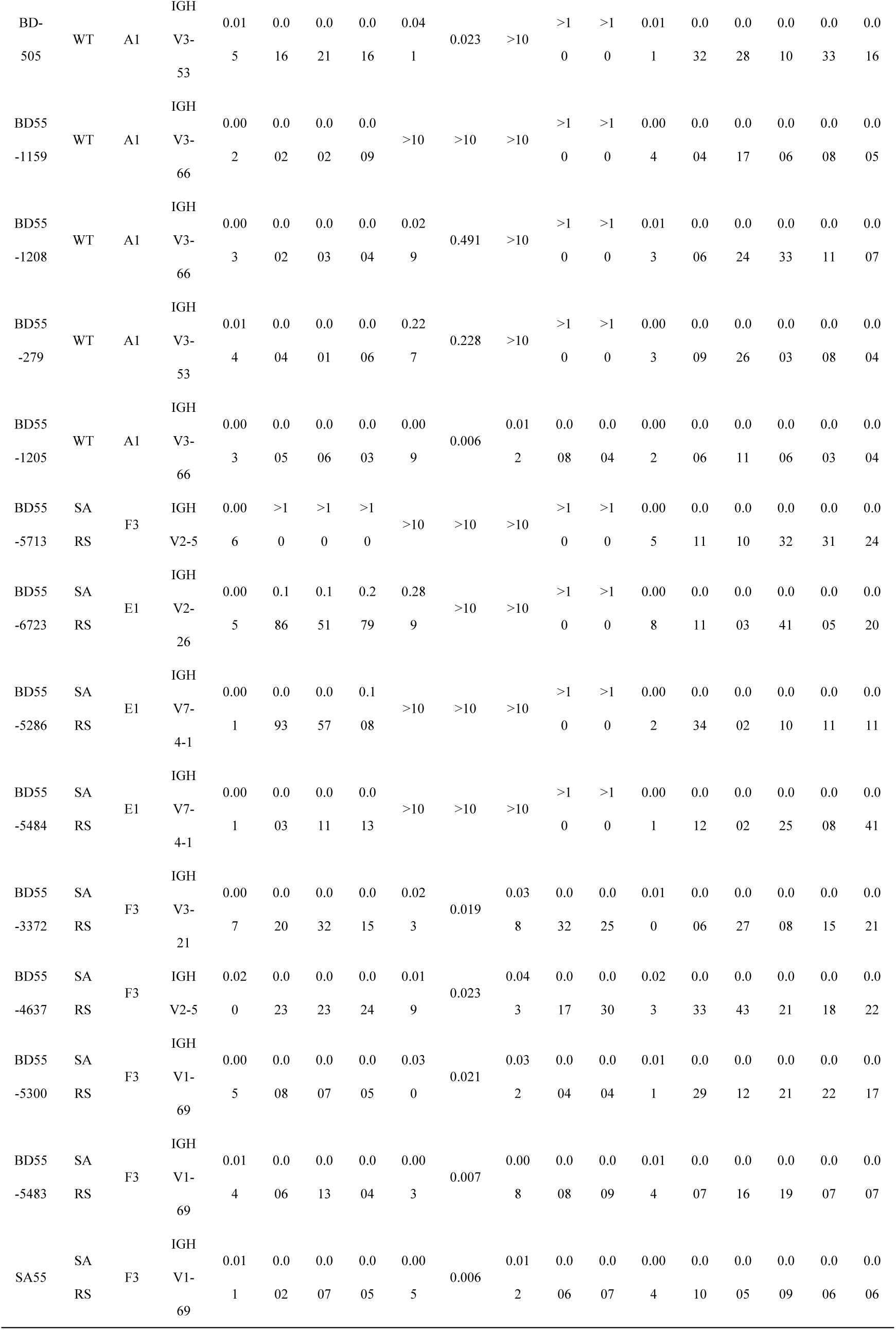
Charcteristics of NAbs isolated from early cohorts that pass the designed filters.

We then evaluated the neutralization against new real-world and prospective variants, including XBB, BA.2.86 or JN.1-derived subvariants with mutations on L455, F456, A475, and Q493 (critical sites targeted by Class 1 antibodies) ^27,36^. BD55-1205 remains potent, whereas many other Class 1 NAbs are escaped by JN.1+F456L and KP.3 (Fig. 3d). Surprisingly, the neutralization of BD55-1205 against KP.3 is even stronger than that against JN.1 and JN.1+F456L, consistent with the stronger ACE2-binding of KP.3 due to F456L-Q493E epistasis, which further underscore the ACE2 mimicry ^36,37^. BD55-1205 IgG demonstrated high apparent affinity to variant RBDs, ranging from 1 pM to 18 nM, in line with its exceptional tolerance to mutations on the epitope (Extended Data Fig. 5c-d).

BD55-1205 also potently inhibited authentic SARS-CoV-2 WT, BA.5.2.1, FL.8, XBB.1.5.6, and JN.3 isolates with IC_50_ from 0.007 to 0.026 μg/mL (Extended Data Fig. 6a). Escape mutation selection with XBB.1.5.6 authentic virus revealed no notable escape in some assay wells even after 12 passages (Extended Data Fig. 6b), and deep sequencing identified only one mutation within BD55-1205 epitope, S490Y (Extended Data Fig. 6c). However, the neutralization activity of BD55-1205 is only slighted reduced against XBB.1.5+S490Y pseudovirus compared to XBB.1.5 (Extended Data Fig. 6d). Interestingly, the ACE2 inhibition efficiency against XBB.1.5+S490Y is also dampened, highlighting the ACE2-mimicking binding again (Extended Data Fig. 6e). Only 8 SARS-CoV-2 sequences with 490Y have been observed from Jan 22 2024 to Jul 19 2024 (https://cov-spectrum.org), indicating low real-world prevalence.

### Structural analyses of BD55-1205

To elucidate the structural basis of BD55-1205’s exceptional breadth, we determined the structure of XBB.1.5 S ectodomain trimer in complex with BD55-1205 Fab using Cryogenic electron microscopy (Cryo-EM) (Extended Data Fig. 7a). We asymmetrically reconstructed the complex structure at an overall resolution of 3.5 Å, with one conformational state referred to as the three “open” RBDs observed (Extended Data Fig. 8a). The high flexibility of three “open” RBDs complicates the alignment during data processing, hence limiting the resolution of the interface in the XBB.1.5 S-BD55-1205 complex. Consequently, we determined the Cryo-EM structure of XBB 1.5 RBD in complex with BD55-1205 Fab (Extended Data Fig. 7b) to further investigate the antibody-RBD interface. The structure of RBD part is similar to a published model from crystallography (Extended Data Fig. 8d).

Consistent with DMS, BD55-1205 is a Class 1 antibody that binds to the apical head of RBD, partially overlapping the RBM and RBD core (Fig. 4a) ^38^. BD55-1205 has a relatively short complementarity-determining region (CDR) H3 of 11 amino acids (IMGT convention) compared to the average in unselected repertoire ^39,40^. The antibody footprint on RBD shows substantial overlap with the hACE2 receptor-binding sites (Fig. 4b and Extended Data Fig. 8b). The distal tip of the RBM deeply inserts into a cavity formed by five CDRs – light chain (LC) CDR1, CDR3 and heavy chain (HC) CDR1-3, resulting in ~1,100 Å^2^ buried area via LC (30%) and HC (70%) contributions (Extended Data Fig. 8c). The epitope of BD55-1205 encompasses over 20 residues, forming an extensive patch along the receptor binding ridge (Extended Data Fig. 8g). RBD:BD55-1205 binding is primarily mediated by polar interactions, many of which involve contacts with RBD carbon backbone. 17 hydrogen bonds are formed between the side chains of RBD residues R403, N405, T415, N417, D420, N487, Y489, Q493, Y501, H505 and HCDR residues Y33, S56, R97, R102, E104, as well as by LCDR residues N30, D93 (Fig. 4c). Additionally, 12 hydrogen bonds are formed between the backbone atoms of 9 RBD residues (L455, R457, K458, Q474, A475, G476, S490, L492, G502) and HCDR (T28, R31, N32, Y33, P53, R102), as well as LCDR (S28) (Fig. 4d-g, Extended Data Fig. 8e-f and 9a). Furthermore, a network of hydrophobic interactions also contributes to the RBD:BD55-1205 interactions, involving G416, Y453, L455, F456 from XBB 1.5 RBD, Y33, Y52, F58, L99, I101 from the HC, as well as W94, P95 from the LC (Fig. 4h). We compared BD55-1205’s interactions with other published Class 1 NAbs that exhibit some extent of neutralization breadth, including P5S-1H1, P5S-2B10, BD-604 and Omi-42 (Fig. 4i-j and Extended Data Fig. 9a-c) ^32,41,42^. Omi-42 is susceptible to A475V and L455F+F456L mutations emergent in several XBB.1.5 and BA.2.86 lineages (Fig. 4i) ^4^. The recognition of P5S-2B10 and P5S-1H1 is affected by N460K, which abrogates the hydrogen bonds on their interfaces (Fig. 4j). However, the extensive interactions with RBD backbone confer BD55-1205’s broad and resilient reactivity (Extended Data Fig. 9a). Despite the unaffected neutralization activity, L455S and A475V moderately dampen the affinity of BD55-1205 to RBD (Extended Data Fig. 5c). This could be explained by the potential impacts on the hydrophobic interactions involving L455, and a hydrogen bond between HC N32 and A475 backbone (Fig. 4h-i). Compared to the other IGHV3-53/3-66-derived NAbs (P5S-2B10, P5S-1H1, BD-604), BD55-1205 has three unique residues in its HCDRs that were introduced by somatic hypermutation (SHM) or VDJ recombination and make contacts with the RBD backbone atoms: R31 on HCDR1, P53 on HCDR2, and R102 on HCDR3 (Extended Data Fig. 9b). These mutations, especially R31 and R102, introduce additional polar interactions at the interface (Fig. 4k). Mutants of BD55-1205 carrying the R31S, P53S, or R102Y substitutions, and the IGHV germline-reverted version (BD55-1205-GLHV) with the mature HCDR3 retained neutralization against BA.5, HK.3.1, JN.1 and JN.1+F456L. However, BD55-1205-GLHV lost neutralizing activity against JN.1+F456L+A475V (Extended Data Fig. 9d), and the apparent affinity of BD55-1205-GLHV to WT, BA.5, XBB.1.5, HK.3 and JN.1 RBD showed a substantial decrease (Extended Data Fig. 9e). These findings reveal that these unique mutations only partially explain BD55-1205’s superior breadth, indicating that the 11-aa HCDR3 is likely the major determinant of the broad reactivity. Nevertheless, these IGHV SHMs enhance its RBD-binding affinity and potentially increase its ability to withstand further antigenic variation.

**Figure 4.**
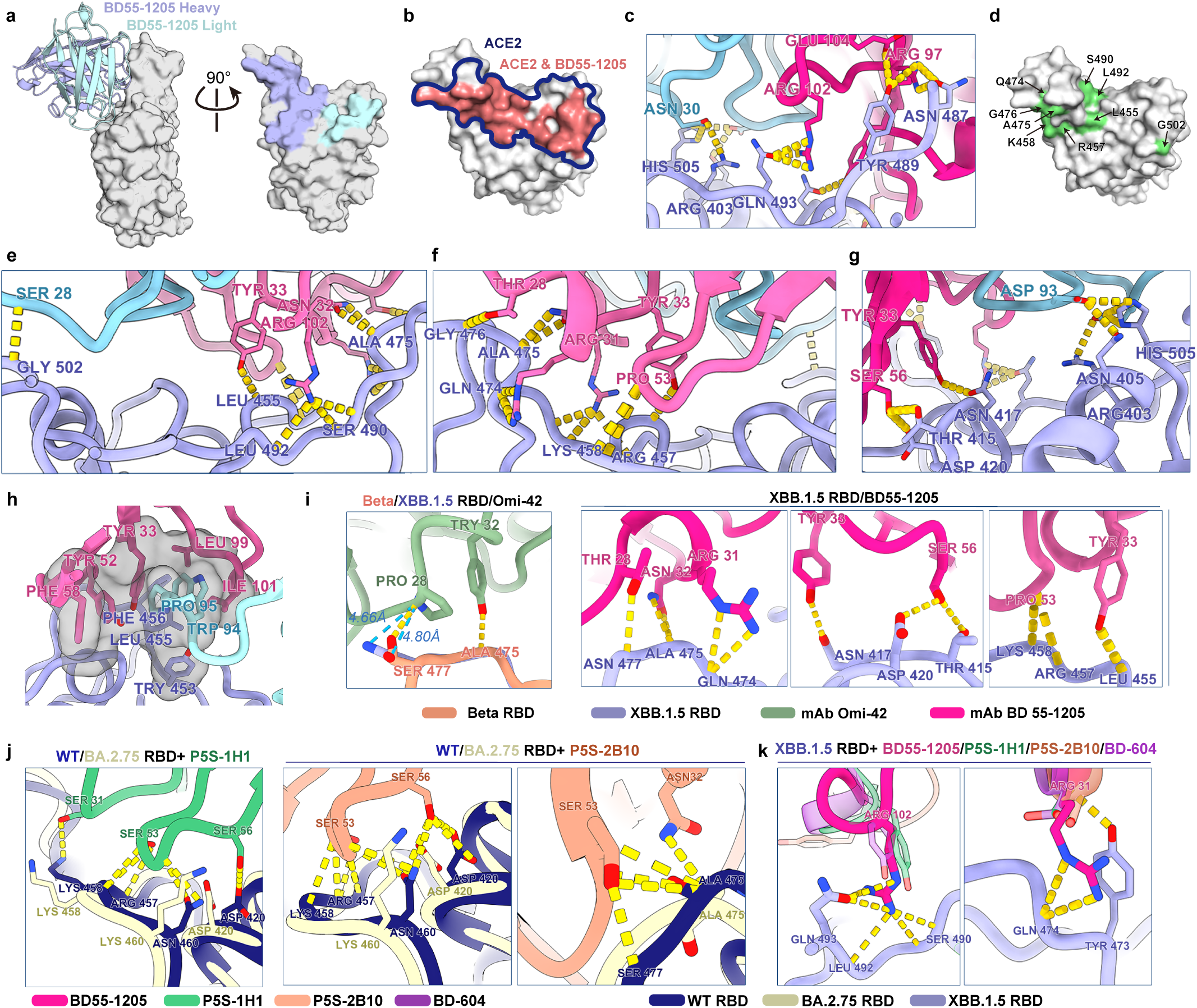
Structural basis for the broad reactivity of BD55-1205. **a**, Structural model of SARS-CoV-2 XBB.1.5 RBD in complex with BD55-1205 from Cryo-EM data. **b**, Overlay of BD55-1205 and hACE2 binding footprints on XBB.1.5 RBD. **c**, Polar interactions between BD55-1205 and XBB.1.5 RBD side chain atoms. **d**, RBD backbone interactions with BD55-1205 Fab. **e-g,** Polar interactions between BD55-1205 heavy chain or light chain and XBB.1.5 RBD backbone atoms in the binding interface. Yellow dashed lines indicate potential polar interactions. RBD, heavy chain, and light chain are colored in blue, magenta, and cyan, respectively. **h**, Hydrophobic interaction between RBD and BD55-1205. **i-k**, Comparison of the RBD interactions with BD55-1205 and other Class 1 NAbs (PDB: Omi-42, 7ZR7; P5S-1H1, 7XS8; P5S-2B10, 7XSC; BD-604, 8HWT).

### mRNA-LNP encoded bnAbs yield high serum neutralizing titers

Recombinant monoclonal antibodies or antibody cocktails have been demonstrated to be clinically active in the prevention of symptomatic COVID ^8–10,43^. Alternative means for delivery of therapeutic antibodies to patients – to the current standard of recombinant mAb products - could prove advantageous towards more rapid and widespread deployment in a pandemic or epidemic context. To that end, we encoded BD55-1205 in mRNA and formulated it into lipid nanoparticles (LNPs) for *in vivo* delivery, similar to mRNA vaccines (see Methods). We introduced the “LA” modification (M428L/N434A) in the Fc region to enhance human FcRn binding at acidic pH, thereby improving the antibody half-life ^44^. Formulated LNPs were delivered by intravenous injection to Tg32-SCID transgenic female mice that are homozygous for human fragment crystallizable neonatal receptor (FcRn) ^45^. To assess BD55-1205 IgG expression kinetics, we quantified human IgG concentration in mouse sera collected at indicated time intervals following mRNA LNP delivery via ELISA(Fig. 5a). We previously reported on in vivo pharmacokinetics of the mRNA-encoded Chikungunya virus E2 glycoprotein specific antibody CHKV-24 (mRNA-1944) determined in a Phase 1 clinical trial ^46,47^. CHKV-24, with a human half-life of 69 days, was employed here as an expression and half-life benchmark and showed expression kinetics highly similar to BD55-1205 in the Tg32-SCID mouse (Fig. 5b) ^46^. At 48 hours, serum antibody levels peaked at 505 μg/mL and 491 μg/mL for BD55-1205 and CHKV-24, respectively (Fig. 5c). Given serum neutralizing titers are an established correlate of protection against SARS-CoV-2 ^43,44,48,49^, we also evaluated the neutralizing titers against XBB.1.5, HK.3.1, and JN.1 pseudoviruses. BD55-1205 achieved high geometric mean peak serum ID_50_ of 4496 against XBB.1.5, 5138 against HK.3.1, and 4608 against JN.1 at 48 hours post administration, underscoring the antibody’s breadth (Fig. 5d-e). As expected, we observed a significant correlation between serum IgG concentrations and ID_50_ against the three variants (Fig. 5f). mRNA-LNP delivery of BD55-1205 to immunocompetent Tg32 mice demonstrated similar peak serum concentrations in both female and male mice (Extended Data Fig. 10a-c). Together, we demonstrate that high serum antibody titers can be achieved following mRNA delivery in a mouse model. The speed and flexibility of mRNA-LNP delivery, coupled with our bnAb prediction methodology, could accelerate the development and deployment of next-generation antibody therapeutics against SARS-CoV-2.

**Figure 5.**
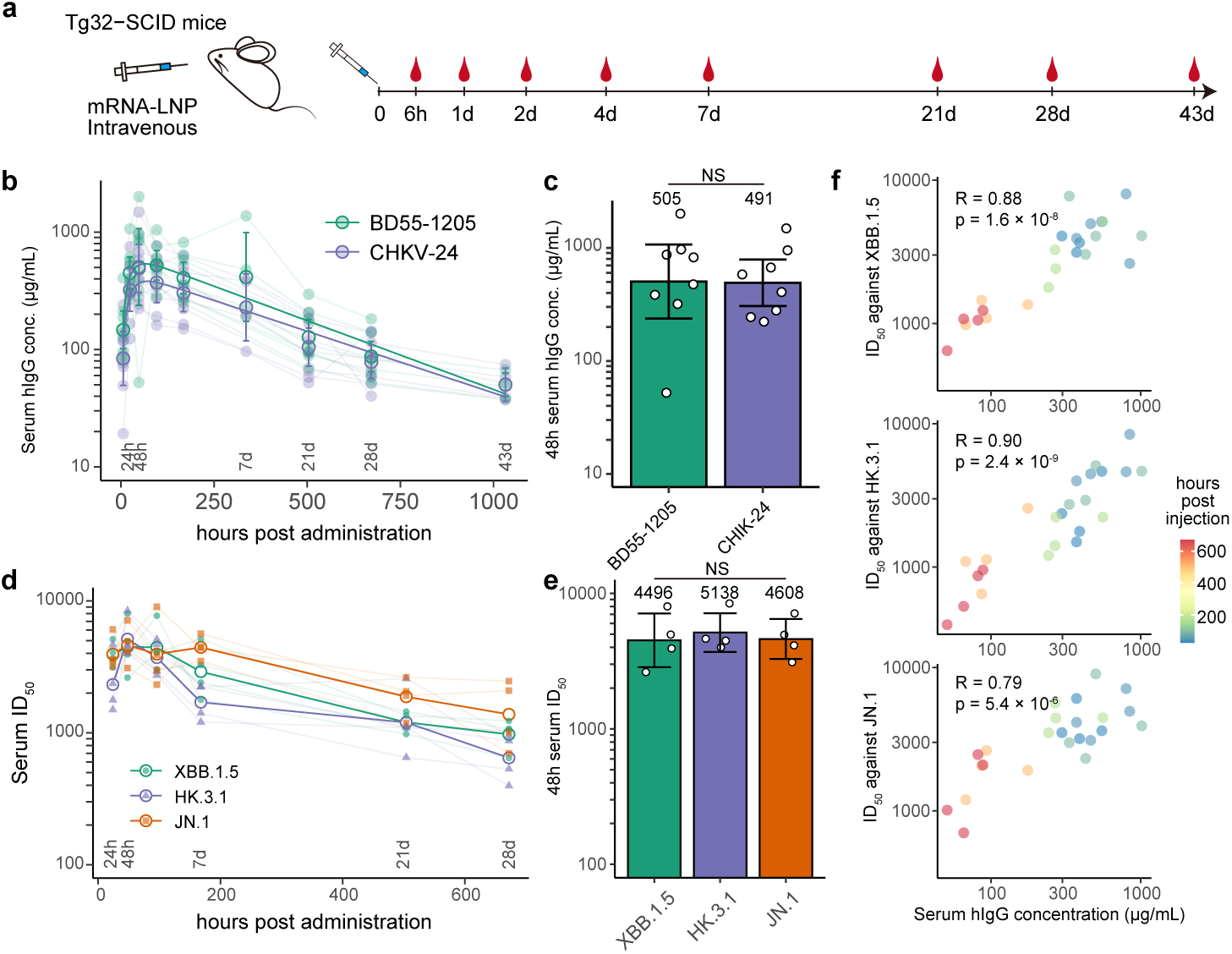
mRNA delivery of BD55-1205 in mice. **a**, A schematic of the experimental design for delivery of BD55-1205 and CHKV-24 via LNP encapsulated mRNA in Tg32-SCID mice. Female mice, 4 per group, received 0.5 mg/kg dose by intravenous injection on day 0 and serum was collected at the indicated time points. **b**, Serum concentration of BD55-1205 and a benchmark antibody CHKV-24 plotted over time. Geometric mean with error (95% confidence interval) is shown by outlined circles with error bars; solid symbols indicate individual animals. A biexponential curve was fitted to the data. Two independent in vivo experiments were combined, each with n=4 animals per group. **c**, Peak serum concentration, occurring at 48 hours post LNP administration, for BD55-1205 (n=8, the two groups are merged) and CHKV-24 (n=4). Bar height and number above the bar indicate the geometric mean; error bars indicate 95% confidence intervals; empty symbols indicate individual animals. NS, not significant (two-sided Wilcoxon rank-sum test). **d**, Half-maximal inhibitory dilutions (ID_50_) of the mouse sera against XBB.1.5, HK.3.1, and JN.1 VSV pseudoviruses for BD55-1205 plotted over time. Geometric mean values are shown as the colored empty circles and lines. The ID_50_ values for individual mouse serum samples are shown as colored points and lines. **e**, Peak serum neutralizing titers in mice receiving BD55-1205 mRNA (n=4, only one group were tested for neutralization) against the three indicated viral variants. Bar height and number above the bar indicate the geometric mean; error bars indicate the 95% confidence interval; empty symbols indicate individual animals. NS, not significant (two-sided Wilcoxon rank-sum test applied to any pair of variants). **f**, Scatter plots showing the correlation between serum hIgG concentrations and the ID_50_ against the three variants at indicated timepoints. Pearson correlation coefficients (R) and the corresponding significance p-values are annotated (two-sided t-test).

## Discussion

In this study, we analyzed a large collection of RBD-targeting mAbs from individuals with diverse SARS-CoV-2 exposure histories to demonstrate the feasibility of high-throughput DMS for accurate and rational variant prediction. This led to the identification of BD55-1205, a Class 1 bnAbs derived from IGHV3-66 in a WT-exposed donor, which neutralized all major variants. While discovered in early 2021, the broad neutralization of BD55-1205 was not appreciated until the emergence of XBB variants in 2023. This underscores the need for rapid identification of such bnAbs and highlights the potential of our prediction framework to facilitate the development of durable antibody drugs early in a pandemic.

Previously, we proposed selecting NAbs targeting immunorecessive epitopes to minimize selection pressure from herd immunity/public Ab responses ^28^. However, this may not apply to newly emergent pathogens or early-pandemic scenarios when epitope immunodominance is unclear. Here we demonstrates that true bnAbs need not exclusively target “rare” epitopes but rather require contacts that can accomodate future mutations. BD55-1205 can withstand high epitope variability over time owing to its high affinity, receptor mimicry and a binding mode that utilizes interactions with RBD backbone atoms. Our study defines key molecular criteria and a platform capable of identifying bnAbs like BD55-1205 in several months, with potential activity retention for multiple years post-discovery.

This rapid bnAb identification process can synergize with the advantages of mRNA-LNP delivery of antibodies. Here, and in past studies, we demonstrated that high peak serum Ab concentrations and neutralizing titers can be achieved with mRNA delivery ^46^. BD55-1205 delivered in mice reached serum levels surpassing CHKV-24, which previously achieved protective levels in clinical trials. Furthermore, the mRNA-LNP platform offers several advantages: (i) rapid product updates during a pandemic, (ii) demonstrated ability to deliver alternative isotypes like IgA for enhanced mucosal protection ^50^, (iii) co-delivery of multiple antibodies to broaden protection against single or multiple pathogens, and (iv) compatibility with multispecific antibodies engineered for superior breadth and potency. This opens up the potential of passive immunotherapy beyond what is possible with conventional antibodies and recombinant Ab delivery.

Although the approach described here offers a promising path to the identification of resilient anti-SARS-CoV-2 mAb therapeutics, it may be constrained by several practical limitations, including: (i) the development of a robust DMS system for new pathogens which may be challenging for some viruses, (ii) the lack established assays to predict efficacy, (iii) insufficient samples for mAb screening, and (iv) unpredictable viral saltations or epistasis beyond current DMS scope.

Altogether, the accurate viral evolution prediction and mRNA-mAb delivery platform described here provide a practical framework for the rapid identification and deployment of bnAbs to combat future SARS-CoV-2 variants. We also envision that this platform could be adapted to respond to other known pathogens with high pandemic potential, such as influenza, or even novel viruses associated with “Disease X” in the future.

## Methods

### Ethic Statement

Authentic SARS-CoV-2 virus strain was undertaken with the approval of the Institutional Biosafety Committee of the Beijing Key Laboratory for Animal Models of Emerging and Remerging Infectious Diseases (Application no. GH2004), and conducted conducted in biosafety level 3 (BSL-3) facility. All animal studies were conducted under an Institutional Animal Care and Use Committee (IACUC)-approved protocol (IACUC 23-07-016) in compliance with the Animal Welfare Act, Public Health Service policy, and other applicable state and city public laws and regulations. Animal studies are designed and executed following guidelines from the Guide for the Care and Use of Laboratory Animals 8th Ed.

### Cell culture

For viral escape assay, African green monkey kidney Vero E6 cells (American Type Culture Collection [ATCC] #CRL-1586/VERO C1008) were maintained in Dulbecco’s Modified Eagle’s Medium (DMEM) high Glucose (Euroclone, Pero, Milan) supplemented with 2 mM L-Glutamine (Euroclone, Pero, Milan), 100 units/mL of penicillin - streptomycin (Gibco, Life Technologies) (“complete DMEM” medium) and 10% Fetal Bovine Serum (FBS) (Euroclone, Pero, Milan).

For plaque reduction neutralization test (PRNT), Vero E6 cells were maintained in MEM (Gibco, Life Technologies) supplemented with 2 mM L-Glutamine, 100 units/mL of penicillin - streptomycin, Non-Essential Amino Acids (Gibco, Life Technologies), HEPES 25mM (Gibco, Life Technologies) and 10% FBS (“PRNT MEM”). Cells were incubated at 37°C in a humidified atmosphere with 5% CO_2_ and passaged every 3 to 4 days.

### Antibody expression and purification

Antibody heavy and light chain genes were codon-optimized and synthesized by GenScript, separately inserted into vector plasmids (pCMV3-CH, pCMV3-CL or pCMV3-CK) via infusion (Vazyme). All antibodies were constructed with human IgG1 constant region. The plasmids were co-transfected into Expi293F cells (ThermoFisher, A14527) using polyethylenimine transfection. The transfected cells were cultured at 36.5°C in 5% CO_2_ for 6-10 days. The expression fluid was collected and centrifuged, and then supernatants containing monoclonal antibodies were purified by Protein A magnetic beads (Genscript, L00695-20). The purified mAb samples were verified by SDS-PAGE (Lablead Biological, P42015).

### Pseudovirus design and neutralization assays

We identify the escape hotspots in SARS-CoV-2 WT exposure cohorts as described previously ^25^. In brief, DMS escape scores of each WT-elicited mAb and each RBD mutation are averaged weighted by the pseudovirus IC_50_ of each mAb against SARS-CoV-2 D614G, and the impacts on ACE2 binding and RBD expression indicated by DMS ^31^.

SARS-CoV-2 wildtype Spike glycoprotein sequence is from the reference genome (MN908947). Spike-pseudotyped viruses of SARS-CoV-2 variants were prepared based on a vesicular stomatitis virus (VSV) pseudovirus packaging system as described previously ^51^. Briefly, variants’ spike plasmid is constructed into pcDNA3.1 vector (BA.1, A67V+HV69-70del+T95I+G142D+V143del+Y144del+Y145del+N211del+L212I+ins214EPE+G339D+S371L+ S373P+S375F+K417N+N440K+G446S+S477N+T478K+E484A+Q493R+G496S+Q498R+N501 Y+Y505H+T547K+D614G+H655Y+N679K+P681H+N764K+D796Y+N856K+Q954H+N969K+L981F; BA.2, T19I+LPPA24-27S+G142D+V213G+G339D+S371F+S373P+S375F+T376A+D405N+R408S+K417N+N440K+ S477N+T478K+E484A+Q493R+Q498R+N501Y+Y505H+D614G+H655Y+N679K+P681H+N7 64K+D796Y+Q954H+N969K; BA.5, BA.2+HV69-70del+L452R+F486V+R493Q; BQ.1.1, BA.5+R346T+K444T+N460K; XBB.1.5, BA.2+V83A+Y144del+H146Q+Q183E+V213E+G339H+R346T+L368I+V445P+G446S+N460K+F486P+F490S+R493Q; HK.3, XBB.1.5+Q52H+L455F+F456L; JN.1, BA.2+ins16MPLF+R21T+S50L+H69-+V70-+V127F+Y144-+F157S+R158G+N211-+L212I+L216F+H245N+A264D+I332V+D339H+K356T+R403K+V445H+G446S+N450D+L45 2W+L455S+N460K+N481K+V483-+A484K+F486P+R493Q+E554K+A570V+P621S+H681R+S939F+P1143L). G*ΔG-VSV (VSV-G pseudotyped virus, Kerafast) and SARS-CoV-2 spike plasmid were transfected to 293T cells (American Type Culture Collection [ATCC], CRL-3216). After culture, the pseudovirus in the supernatant was harvested, filtered, aliquoted, and frozen at −80°C for further use.

Huh-7 cell line (Japanese Collection of Research Bioresources [JCRB], 0403) was used in pseudovirus neutralization assays. Purified mAbs were serially diluted in culture media and mixed with pseudovirus, and incubated for 1 h in a 37°C incubator with 5% CO_2_. Then, the digested Huh-7 cells were seeded in the antibody-virus mixture. After one day of culture in the incubator, the supernatant was discarded, and D-luciferin reagent (Vazyme, DD1209-03) was added into the plates. After 2 min incubation in darkness, cell lysis was transferred to detection plates. The luminescence values were detected and recorded with a microplate spectrophotometer (PerkinElmer, HH3400). IC_50_ was determined by fitting four-parameter logistic regression models.

### Authentic virus propagation and titration

Authentic SARS-CoV-2 Omicron variant XBB.1.5.6 was kindly provided by Rega Institute Leuven (Belgium). Live Omicron variant BA.2.86 sub-lineage JN.3 was kindly provided by the Medicines and Healthcare Product Regulatory Agency (MHRA, United Kingdom).

Viral propagation of both strains was carried out as previously described ^52^. Briefly, 175cm^2^ flasks were inoculated with Vero E6 cells diluted in complete DMEM 2% FBS (1×10^6^ cells/mL). Cells were incubated at 37°C, 5% CO_2_ in a humidified atmosphere for 20-24 hours, then the sub-confluent cell monolayer was washed twice with sterile Dulbecco’s phosphate buffered saline (DPBS) and inoculated with the SARS-CoV-2 virus at a multiplicity of infection (MOI) of 0.001. After 1 hour at 37°C, 5% CO_2_, the flasks were filled with 50 mL of complete DMEM 2% FBS and kept at 37°C, 5% CO_2_. Flasks were inspected daily under an optical microscope to check for signs of cytopathic effect (CPE) in the Vero E6 monolayer. Once CPE was developed in at least 80-90% of the cell monolayer, the supernatants of the infected cell culture were collected, centrifuged at 1000 rpm (106 ×g) for 5 minutes (4°C), aliquoted and stored at −80°C.

For viral escape assay, a titration of the propagated Omicron variant XBB.1.5.6 was performed in 96-well plates containing confluent Vero E6 cells, by means of the 50% tissue culture infectious dose assay (TCID_50_). Cells infected with serial 10-fold dilutions of the virus (from 10^−1^ to 10^−11^) were incubated at 37°C, 5% CO_2_ and monitored for signs of virus-induced CPE under an inverted optical microscope for 4 days. The end-point viral titer, defined as the reciprocal of the highest viral dilution resulting in at least 50% CPE in the inoculated wells, was calculated according to the Reed and Muench formula ^53^. Titration of plaque forming unit/ml (PFU/ml) of SARS-CoV-2 Omicron XBB.1.5.6 and JN.3 variants was performed in pre-seeded Vero E6 cells in 96-well plates. Briefly, cells were infected with serial 0.5 Log-fold dilutions of the virus (from 10^−1^ to 10^−6^) and incubated for 24 h at 37°C, 5% CO_2_. The viral titer was calculated by PFU counting.

### Authentic virus escape mutant escape assay

For the authentic virus escape assay, an experimentally determined concentration of authentic XBB.1.5.6 virus was sequentially passaged in Vero E6 cells in the presence of serially diluted SARS-CoV-2-specific-monoclonal antibody (BD55-1205). The viral escape assay was performed as previously reported ^52^. Briefly, 12 serial two-fold dilutions of each antibody sample were prepared in DMEM 2% FBS (starting concentration of antibody before virus addition: 20 μg/mL). Each serially diluted sample was added to the wells of a 24-well plate, pre-seeded with Vero E6 cells (2×10^5^ cells/well). Then, a virus solution containing 10^5^ TCID_50_ of authentic SARS-CoV-2 Omicron variant XBB.1.5.6 was dispensed in each antibody-containing well, and in wells dedicated to virus-only control.

The plates were then incubated for 1 h at 37°C, 5% CO_2_, to allow binding of the antibody sample to the virus. The virus-sample mixture was then transferred into the wells of a 24-well plate containing previously seeded Vero E6 cells, to allow their infection from the unbounded residual virus. The plates were incubated for 7 days at 37°C, 5% CO_2_, then cells were examined for the presence of virus-induced CPE using an inverted optical microscope. The content of the well corresponding to the highest antibody concentration showing complete CPE was collected and further diluted to be used as the viral solution in the next virus passage. The potency of each antibody was recorded at each virus passage and expressed as Inhibitory Concentration 100% (IC100) (i.e., the lowest antibody concentration inhibiting development of CPE). The virus pressured with SARS-CoV-2 antibody was passaged in cell cultures along with the antibody sample of interest until CPE was observed at higher antibody concentrations. At each passage, both the virus pressured with the antibody sample of interest and the virus-only control were harvested, propagated for one round of passaging in different 25cm^2^ flasks (pre-seeded with 1×10^5^ Vero E6 cells/mL), aliquoted and stored at −80°C to be used for RNA extraction and sequencing. The sequences of both these types of samples can assist in distinguishing between adaptation to cell culture conditions and escape mutations. Parallel titrations of each antibody-pressured virus were performed at every passage in 96-well plates containing pre-seeded Vero E6 cells (1.5×10^4^ cells/well), to monitor the viral titer at each test.

### RNA Extraction

The RNA extraction to obtain the viral genetic material for next-generation sequencing was performed using biocomma® Nucleic Acid Purification Kit (Spin Column) commercial kit (MNP027-1E, Biocomma Limited), as described previously ^52^. Briefly, 300 μL of viral sample were mixed with 500 μL of “Buffer GLX”, vortexed for 1 minute and incubated at room temperature (RT) for at least 5 minutes to allow virus lysis. The supernatant was then transferred into a spin column inserted in a collection tube and centrifuged at 12,000 rpm (15,287×g) for 1 minute at RT. After discarding the flow-through, 500 μL of “Buffer PD” (previously re-suspended with isopropanol) were added, centrifuged as before, followed by elimination of the eluted solution. The column was then washed with 700 μL of “Buffer PW” (previously re-suspended in absolute ethanol), centrifuged and eluted as before. This step was repeated twice. The spin column was then centrifuged at 12,000 rpm (15,287×g) for 2 minutes and left with open lid for 5 minutes to allow evaporation of residual ethanol. The column was placed in a new collection tube and 60 μL of RNAse-free ddH20 were added. After a 2 minutes incubation at RT, the column was centrifuged for 2 minutes at 12,000 rpm (15,287×g) to elute and collect the RNA, which was stored at −80°C until shipment for sequencing.

### Deep sequencing of authenic virus from escape assay and data analysis

The cDNA preparation was performed in a total volume of 40 uL by following manufacturer’s recommendations for SuperScript™ II Reverse Transcriptase (Life Technologies 18064022), Random Hexamers (50 µM) (Euroclone N8080127) and RNaseOUT™ Recombinant Ribonuclease Inhibitor (Life Technologies 10777019) suing a thermocycler. Sars-CoV-2 genome amplicons were generated using ARTIC v3.5.2 panel (IDT, cat#10016495), together with a set of custom oligo pools to improve coverage on the RBD domain sequence. The Celero™ DNA-Seq kit (NuGEN, San Carlos, CA, USA) was then used for library preparation following the manufacturer’s instructions. Both input and final libraries were quantified by Qubit 2.0 Fluorometer (Invitrogen, Carlsbad, CA, USA) and quality tested by Agilent 2100 Bioanalyzer High Sensitivity DNA assay (Agilent technologies, Santa Clara, CA, USA). Libraries were then prepared for sequencing and sequenced on Illumina NovaSeq6000 (Illumina, San Diego, CA, USA) in paired-end 150 mode to generate a minimum of 5 million reads per sample.

Raw sequencing reads were first processed by removing PCR primer sequences using Cutadapt v2.6, with parameters set to discard untrimmed pairs and ensure a minimum read length of 50 bp. Quality trimming was performed with Fastp v0.20.0 to retain bases with a Phred score ≥ Q30. The resulting high-quality reads were aligned to the NCBI Reference Sequence accession “OQ063792.1”. Subsequent filtering wiat a custom script removed reads with low mapping quality or suboptimal alignment characteristics. Variants were identified using GATK HaplotypeCaller v4.1.6.0 and normalized with Bcftools norm v1.9. A consensus sequence for each sample was generated by applying filtered variants to the reference sequence using Bcftools consensus, with low-coverage positions masked. Coverage was assessed with Bedtools genomecov v2.29.2.

### Authentic virus plaque reduction neutralization

Determination of half maximal inhibitory concentration (IC_50_) was performed in Vero E6 cells by immunodetection of viral antigen. Briefly, 100 PFU/well of SARS-CoV-2 Omicron XBB.1.5.6 or Omicron JN.3 were incubated 1h with 5% CO_2_ with serial 4-fold dilutions of monoclonal antibodies (range 50-0.003 nM). At the end of incubation, pre-seeded Vero E6 cells in 96-well plates were adsorbed with virus-sample mixture for 1 h at 37 °C with 5% CO_2_. After removal of virus inoculum, the overlay media was added at each well and plates were incubated for 24 h at 37 °C with 5% CO_2_. The immunodetection assay was performed as described previously ^54^ with minor modifications. Briefly, cells were fixed for 3 hours with formalin 10% (Sigma Aldrich, HT501320), and permeabilized for 20 min with 0.1% Triton X-100 (Sigma Aldrich, 1.08603). After washing with PBS 1X (ThermoFisher, 28348) containing 0.05% Tween 20 (Sigma Aldrich), plates were incubated for 1 h with SARS-CoV-2 nucleocapsid mouse monoclonal antibody (Genscript, A02048) diluted 1:1000 in blocking buffer (PBS 1X containing 1% BSA (Sigma Aldrich, A8327) and 0.1% Tween 20). After washing, cells were incubated for 1 h with a polyclonal Horseradish Peroxidase (HRP)-coupled anti-mouse IgG secondary antibody (ThermoFisher, 31430) diluted 1:2000 in blocking buffer. Next, cells were washed and the TrueBlue™ Peroxidase Substrate (Sera Care, 5510-0030) was added to each well.

Detection of microplaques was performed with ImmunoSpot® S6 Ultra-V Analyzer (C.T.L.) reader using BioSpot® software according to instrument specifications. The IC_50_ for each sample was calculated with GraphPad Prism software (v9.0.1) using the dose-response inhibition category and apply log(inhibitor) vs. normalized response - Variable slope.

Neutralization assays against SARS-CoV-2 WT (Wuhan-Hu-1), BA.1 (EPI_ISL_8187354), BA.5.2.1 (EPI_ISL_17261619.2), and FL.8 (XBB.1.9.1.8, EPI_ISL_17262369) authentic virus were performed in a different lab. In brief, purified BD55-1205 is subsequently diluted in two-fold from 500 ng/mL to 0.244 ng/mL. These diluted antibodies were mixed with live virus suspension containing 100 TCID_50_ and added to 96-well plates at a 1:1 ratio. The plates were incubated in a 36.5 °C incubator with 5% CO_2_ for 2 h. Following the incubation, Vero cells (a gift from WHO, (ATCC, CCL-81)) were added to each well containing the antibody-virus mixture. The plates were further incubated for 5 days in an incubator with 5% CO_2_ at 36.5 °C. Cytopathic effects were evaluated by microscopy, and the IC_50_ values were determined by fitting two-parameter Hill equations. Experiments were conducted in four biological replicates in a biosafety level 3 (BSL-3) facility.

### Preparation of spike-pseudotyped rVSV

Similar to previous reports, SARS-CoV-2 XBB.1.5 S-pseudotyped rVSV was constructed and rescued from DNA clones ^55,56^. In brief, the VSV G gene on the plasmid encoding VSV anti-genome with T7 promoter was replaced by codon-optimized SARS-CoV-2 XBB.1.5 Spike gene with a C-terminal 21-aa deletion. GFP reporter gene was inserted into the VSV genome before the Spike gene. BHK21 cells were infected with the vaccinia virus vTF7-3 that expresses T7 polymerase for 2 hours, and the supernatant was discarded. The VSV antigenome plasmid, and helper plasmids encoding VSV N, P, G, and L genes (Kerafast, EH1013-1016) were co-transfected to the cells using Lipofectamine 3000. After 48-hour incubation, the supernatant was collected, filtered by a 0.22 μm filter, and passaged in Vero E6 cells. The virus was passaged every 2-3 days. After 3-4 times amplification, the viral RNA was extracted and amplified by reverse transcription PCR. The Spike gene was then amplified and sequenced for validation. The supernatant that contains the rescued virus was aliquoted and stored at −80°C.

### rVSV-based escape mutation screening under antibody pressure

Monoclonal antibodies were prepared at concentrations of 5, 1.25, 0.3125, and 0.078 µg/mL in Dulbecco’s Modified Eagle Medium (Hyclone, SH30243.01) supplemented with 2% fetal bovine serum (Hyclone, SH30406.05). In some replicates, an additional concentration of 20 µg/mL was also included. A volume of 0.5 mL of each dilution was added to individual wells of a 24-well plate. Subsequently, 0.5 mL of XBB.1.5-S-pseudotyped rVSV with a titer of 4×10^6^ focus-forming units per mL (FFU/mL) was introduced to each well, resulting in a total volume of 1 mL per well. The plates were incubated at room temperature for 30 minutes to allow antibody-virus binding.

After incubation, 200 µL of Vero cell suspension (1×10^6^ cells/mL) was added to each well, bringing the final volume to 1.2 mL. The cell-virus-antibody mixture was then cultured at 37°C in an atmosphere containing 5% CO_2_ for 72 hours. The fluorescence signal, indicating successful viral entry and replication, was captured using a BioTek Cytation 5 Cell Imaging Multimode Reader (Agilent).

For the viral passage experiment, the supernatants from GFP-positive wells that contained the highest concentration of antibodies were harvested. The samples were centrifuged at 350 g for 3 minutes, and the clarified supernatants were then diluted in DMEM supplemented with 2% FBS, and mixed with antibodies at various concentrations and incubated for 30 minutes before the addition of Vero cells. Subsequent passages were performed under identical conditions to those of the initial experiment, until the virus could successfully infected cells (GFP^+^) under the pressure of antibody at the highest concentration.

### Recombinant RBD expression and purification

DNA fragments that encode SARS-CoV-2 variant RBD (Spike 319-541) were codon-optimized for human cell expression and synthesized by Genscript. His-AVI tags were added at the end of the RBD gene fragments. The fragments were then inserted into pCMV3 vectors through infusion (Vazyme). The recombination products were transformed into E. coli DH5α competent cells (Tsingke). Colonies with the desired plasmids were confirmed by Sanger sequencing (Azenta) and cultured for plasmid extraction (CWBIO). Expi293F cells were transfected with the constructed plasmids and cultured for 6 days. The products were purified using Ni-NTA columns (Changzhou Smart-lifesciences, SA005100) and the purified samples were verified by SDS-PAGE.

### Surface plasmon resonance

SPR experiments were performed on Biacore 8K (Cytiva). mAbs (human IgG1) were immobilized onto Protein A sensor chips (Cytiva, 29127556). Purified SARS-CoV-2 variant RBDs were prepared in serial dilutions (1.5625, 3.125, 6.25, 12.5, 25, and 50 nM) and injected over the sensor chips. The response units were recorded by Biacore 8K Control Software (Cytiva, v4.0.8.19879) at room temperature, and the raw data curves were fitted to a 1:1 binding model to determine the affinities (K_D_) using Biacore 8K Evaluation Software (Cytiva, v4.0.8.20368).

### Protein expression and purification for Cryo-EM

The Spike gene of XBB.1.5 (T19I, Δ24-26, A27S, V83A, G142D, Δ144, H146Q, Q183E, V213E, G252V, G339H, R346T, L368I, S371F, S373P, S375F, T376A, D405N, R408S, K417N, N440K, V445P, G446S, N460K, S477N, T478K, E484A, F486P, F490S, R493Q, Q498R, N501Y, Y505H, D614G, H655Y, N679K, P681H, N764K, D796Y, Q954H, N969K) and XBB.1.5 RBD were realized by overlapping PCR with the full-length S gene (residues 1-1208, GenBank: MN908947) as template. The S gene was constructed into the vector pCAGGS with a T4 fibrin trimerization motif and a HRV3C protease site and a Twin-Strep-tag at the C-terminal of spike and RBD sequences to facilitate protein purification and was mutated as previously described ^57^. All the constructed vector were transiently transfected into suspended FreeStyle^TM^ 293-F cells (ThermoFisher, R79007) and cultured at 37 °C in a rotating, humidified incubator supplied with 8% CO_2_ and maintained at 130 rpm (0.25×g). After incubation for 72h, the supernatant was harvested, concentrated, and exchanged into the binding buffer by tangential flow filtration cassette. The S proteins were then separated by chromatography using resin attached with streptavidin and further purified by size exclusion chromatography using a Superose 6 10/300(GE Healthcare) in 20 mM Tris, 200mM NaCl, pH 8.0.

### Production of Fab fragment

To generate the Fab fragments for Cryo-EM analyses, the purified antibodies were processed using the Pierce Fab preparation kit (ThermoFisher, 44895) as described previously ^13^. Briefly, the samples were first applied to desalination columns to remove the salt. After centrifugation, the flow through was collected and incubated with beads attached with papain to cleave Fab fragments from the whole antibodies. Then the mixtures were transferred to Protein A affinity column which specifically binds the Fc fragments of antibodies. After centrifugation, the Fab fragments were obtained and dialyzed into Phosphate Buffered Saline (PBS).

### Cryo-EM sample collection, data acquisition and structure determination

The cryo-EM samples of S trimers in complex with BD55-1205 with a molar ratio of 1:4 (S protein:BD55-1205) on ice to obtain S-BD55-1205 complex. Then, the complex was deposited onto the freshly glow-discharged grids (C-flat 1.2/1.3 Au). After 6 seconds’ blotting in 100% relative humidity, the grid was plunged into liquid ethane automatically by Vitrobot (FEI). Cryo-EM data sets were collected at a 200 kV FEI Talos Arctica microscope equipped with a K2 detector. Movies (32 frames, each 0.2 s, total dose of 60 e/Å^−2^) were recorded with a defocus range between 1.5-2.7 μm. Automated single particle data acquisition was carried out by SerialEM, with a calibrated magnification of 75,000, yielding a final pixel size of 1.04 Å. A total of 5722 micrographs were collected. CryoSPARC (v4.4.1) was used to correct beam induced motion and average frames. Then, the defocus value of each micrograph was estimated by patch CTF estimation. 2,515,383 particles of XBB.1.5 S-BD55-1205 complex were autopicked and extracted for further 2D classification and hetero-refinement. After that, 238,788 particles of XBB.1.5 S-BD55-1205 complex were used for homo-refinement in cryoSPARC for the final cryo-EM density.

To improve the resolution of the binding surface of RBD-antibody, the cryo-EM sample of XBB.1.5 RBD in complex with BD55-1205 and BD57-0120 Fab, which is another RBD-targeting mAb that doesn’t compete with BD55-1205, was also deposited. This approach allowed us to deduce a more accurate epitope and paratope than was achievable using the flexible up RBD conformation in the BD55-1205-S cryo-EM structure, with BD57-0120 Fab utilized to increase the molecular weight of the complex. We performed asymmetric reconstruction of the complex structure, achieving an overall resolution of 3.3 Å, enabling reliable analysis of the interaction interface.The cryo-EM samples of XBB.1.5 RBD in complex with BD55-1205 were mixed in a molar ratio of 1:1.2:1.2 (RDB: BD55-1205: BD57-0120). Movies (32 frames, each 0.2 s, total dose of 60 e /Å^−2^) were recorded using a Falcon 4 Summit direct detector with a defocus range between 1.2-1.8 μm. Automated single particle data acquisition was carried out by EPU, with a calibrated magnification of 96,000, yielding a final pixel size of 0.808 Å. A total of 5,866 micrographs for XBB.1.5 RBD-BD55-1205/BD57-0120 complex were collected. CryoSPARC was used to correct beam induced motion and average frames. Then, the defocus value of each micrograph was estimated by patch CTF estimation. 3,186,044 particles of XBB.1.5 RBD-BD55-1205/BD57-0120 complex were autopicked and extracted for further 2D classification and hetero-refinement. After that, 266,321 particles of XBB.1.5 RBD-BD55-1205/BD57-0120 complex were used for homo-refinement and non-uniform refinement in cryoSPARC for the final cryo-EM density.

The resolutions were evaluated on the basis of the gold-standard Fourier shell correction (threshold = 0.143) and evaluated by ResMap (v1.1.4). All dataset processing is shown in the Extended Data Fig. 6.

### Structure model fitting and refinement

The atom models of the complex were first fitting the chain of the apo (PDB: 7XNQ) and Fab (predicted by AlphaFold) into the obtained cryo-EM density by Chimera (v1.5). Then the structure was manually adjusted and corrected according to the protein sequences and density in Coot (v0.9.8.91), real-space refinement was performed by Phenix (v1.20.1).

### Generation of modified mRNA and LNPs

BD55-1205 with the ‘LA’ modification and CHKV-24 with the ‘LS” modification (for consistency with histrocial data) were used in the studies. Sequence-optimized mRNA encoding functional IgG monoclonal antibodies were synthesized *in vitro* using an optimized T7 RNA polymerase-mediated transcription reaction with complete replacement of uridine by N1-methyl-pseudouridine ^58^. The reactions included a DNA template containing an open reading frame flanked by 5′ untranslated region (UTR) and 3′ UTR sequences with a terminal encoded polyA tail. Free mRNA was purified, buffer exchanged and sterile filtered.

Lipid nanoparticle-formulated mRNA was produced through a modified ethanol-drop nanoprecipitation process as described previously ^47,50,59–62^. Briefly, ionizable, structural, helper, and polyethylene glycol lipids were mixed with mRNA in acetate buffe at a pH of 5.0 and at a ratio of 3:1 (lipids:mRNA); 2:1 ratio Heavy Chain to Light Chain. The mixture was neutralized with Tris-Cl at a pH 7.5, sucrose was added as a cryoprotectant, and the final solution was sterile filtered. Vials were filled with formulated LNP and stored frozen at –70°C until further use. All formulations underwent analytical characterization for particle size, RNA encapsulation, mRNA purity, osmolarity and endotoxin and were found to be between 80 to 100 nm in size, with greater than 90% encapsulation and < 1 EU/ml of endotoxin, and the material was deemed acceptable for *in vivo* study.

Moderna’s proprietary UTR sequences for enhancing mRNA stability and translation efficiency are protected under U.S. Patent No. 11389546, and the methods for LNP composition and delivery are protected under U.S. Patent No. 11285222.

### Expression of mAbs in mice

Homozygous ‘Tg32-SCID’ mice (B6.Cg-Fcgrt^tm1Dcr^ Prkdcscid Tg(FCGRT)32Dcr/DcrJ, cat. no. 018441) and ‘Tg32’ mice (B6.Cg-Fcgrt^tm1Dcr^ Tg(FCGRT)32Dcr/Dry, cat. no. 014565) were obtained from The Jackson Laboratory, Bar Harbour, Maine. Animals were housed in groups of 4, fed standard chow diets, subjected to a photoperiod of 12 hours on, 12 hours off dark/light cycle and kept at an ambient animal room temperature of 70° ± 2° F with a room humidity of 50% ± 5%.

Six- to eight-week-old female Tg32-SCID mice (2 groups, 4 mice in each group) or male (1 group with 4 mice) and female (1 group with 4 mice) Tg32 mice (The Jackson Laboratory) were injected intravenously with the indicated mRNA LNP, at the indicated dose (0.5 mg/kg) in 100 µL total volume. All mRNA LNPs for *in vivo* were prepared by coformulations of HC and LC for the expression of the full IgG. Mice were bled via submandibular vein at the indicated time points and serum was isolated for antibody quantification by ELISA and assessment of serum neutralizing titer. At the final indicated time point, mice were euthanized via CO_2_ asphyxiation and a terminal bleed was collected via cardiac puncture.

### Mouse serum ELISA for human IgG quantification

To quantitate total human IgG (hIgG), 96-well NUNC Maxisorp plates (ThermoScientific, 439454) were coated with 0.1 mL per well of goat anti-human IgG Fc fragment (Bethyl A80-104A) at 1:100 dilution in 0.05 M carbonate-bicarbonate (Sigma C30411) overnight at 4°C. Plates were washed with an automated plate washer (Biotek) 4x with 0.05% PBS-T and were subsequently blocked with 0.2 mL per well of Superblock PBST (ThermoFisher, 37515) for 1.5 hour at 37°C. Using purified antibodies as a standard, mRNA transfection supernatant or mouse serum was serially diluted in PBST in a dilution plate and 0.1mL per well was transferred to the coated plates and incubated for 2 hour at 37°C. Following incubation, plates were subsequently washed and incubated with 0.1 mL per well of goat anti-human IgG horseradish peroxidase (HRP; Southern Biotech; 2040-05, 1:5000), for 1 hour at 37°C. Plates were subsequently washed and incubated with 0.1 mL per well of Sureblue TMB 1-C substrate (Sera care, 52-00-04) for 10 minutes. The reaction was stopped with 0.1 mL per well of TMB Stop solution (SeraCare, 50-85-05) and read at an absorbance of 450nm on a SpectraMax ABS Microplate Reader (Molecular Devices). Absolute quantities of human antibody in transfection supernatant or mouse serum were extrapolated in GraphPad Prism (v9.0.1) using a standard curve.

### Mice serum pseudovirus neutralization assays

Codon-optimized full-length spike genes (XBB.1.5, JN.1 and HK.3.1) were cloned into a pCAGGS vector. Spike genes contained the following mutations: (a) XBB.1.5: T19I, L24-, P25-, P26-, A27S, V83A, G142D, Y144-, H146Q, Q183E, V213E, G252V, G339H, R346T, L368I, S371F, S373P, S375F, T376A, D405N, R408S, K417N, N440K, V445P, G446S, N460K, S477N, T478K, E484A, F486P, F490S, Q498R, N501Y, Y505H, D614G, H655Y, N679K, P681H, N764K, D796Y, Q954H, N969K; (b) JN.1: ins16MPLF, T19I, R21T, L24-, P25-, P26-, A27S, S50L, H69-, V70-, V127F, G142D, Y144-, F157S, R158G, N211-, L212I, V213G, L216F, H245N, A264D, I332V, G339H, K356T, S371F, S373P, S375F, T376A, R403K, D405N, R408S, K417N, N440K, V445H, G446S, N450D, L452W, L455S, N460K, S477N, T478K, N481K, V483-, E484K, F486P, Q498R, N501Y, Y505H, E554K, A570V, D614G, P621S, H655Y, N679K, P681R, N764K, D796Y, S939F, Q954H, N969K, P1143L; and (c) HK.3.1: T19I, L24-, P25-, P26-, A27S, Q52H, V83A, G142D, Y144-, H146Q, F157L, Q183E, V213E, G252V, G339H, R346T, L368I, S371F, S373P, S375F, T376A, D405N, R408S, K417N, N440K, V445P, G446S, L455F, F456L, N460K, S477N, T478K, E484A, F486P, F490S, Q498R, N501Y, Y505H, D614G, H655Y, N679K, P681H, N764K, D796Y, Q954H, N969K. To generate VSVΔG-based SARS-CoV-2 pseudovirus, BHK-21/WI-2 cells were transfected with the spike expression plasmid and infected by VSVΔG-firefly-luciferase as previously described ^63^. VeroE6 cells were used as target cells for the neutralization assay and maintained in DMEM supplemented with 10% FBS. To perform neutralization assay, mouse serum samples were heat-inactivated for 45 minutes at 56 °C, and serial dilutions were made in DMEM supplemented with 10% FBS. The diluted serum samples or culture medium (serving as virus-only control) were mixed with VSVΔG-based SARS-CoV-2 pseudovirus and incubated at 37 °C for 45 minutes. The inoculum virus or virus–serum mix was subsequently used to infect VeroE6 cells (ATCC, CRL-1586) for 18 hours at 37 °C. At 18 hours after infection, an equal volume of One-Glo reagent (Promega, E6120) was added to culture medium for readout using a BMG PHERastar-FSX plate reader. The percentage of neutralization was calculated based on relative light units (RLUs) of the virus control and subsequently analyzed using four-parameter logistic curve (GraphPad Prism 8.0).

### Statistics & Repreducibility

The sample sizes in this study (number of experimental measurement replicates and number of animals) are based on experience and sufficient for necessary statistical tests. Experimental assays were performed in at least two independent experiments. No statistical method was used to predetermine sample size. 668 mAbs were excluded from the study because of insufficient amount of antibody and missing neutralization data against D614G or autologous pseudovirus. No data were excluded from the analyses. Results of all replicates are shown in the figure. The experiments were not randomized. The investigators were not blinded to allocation during experiments and outcome assessments.

In Fig. 2e, enrichment of Omicron-effective NAbs among NAbs neutralizing designed mutants is indicated by hypergeometric tests of the number of mAbs with IC_50_ against a designed mutant < 0.05 μg/mL, and the number of mAbs with IC_50_ against a real-world Omicron strain < 0.05 μg/mL. In Fig. 5c and 5e, we used Wilcoxon rank-sum tests to determine whether there are significant differences between mice groups, where NS indicates not significant. Non-parametric Wilcoxon tests do not assume the data normality. For t-tests involved in the calculation of Pearson’s correlation, data distribution was assumed to be normal but this was not formally tested. Illustration was performed by PyMOL (v2.6.0a0), Python package logomaker (v0.8), R package ggplot2 (v3.4.4) and ComplexHeatmap (v2.14).

## Data availability

Information of the monoclonal antibodies involved in this study is included in Table S1. Cryo-EM data for structures have been deposited in the Protein Data Bank (PDB) with accession 8XE9 and 8XEA, and in the Electron Microscopy Data Bank (EMDB) with accession EMD-38283 and EMD-38284. Other necessary source data to reproduce the analyses and illustrations have been deposited to Zenedo (10.5281/zenodo.15294998). Additional materials and data are available from the lead corresponding author (Y.C.) upon request and subject to a Material and Data Transfer Agreement. All inquiries will be replied within 7 working days.

## Code availability

Custom scripts for data analysis and illustrations involved in this study have been deposited to GitHub (https://github.com/yunlongcaolab/predict-identify-bnabs).

## Acknowledgments

This project is financially supported by the Ministry of Science and Technology of China (2022ZD0115002, Y.C.) and National Natural Science Foundation of China (32222030, Y.C.).

We wish to acknowledge Giulia Piccini, Alessandro Manenti, Emanuele Montomoli, all at VisMederi Srl, Sienna, Italy and their lab team, and Zhe Lv at Sinovac Inc., for their contribution of methods and the performance of authentic virus assays. We also thank the Medicines and Healthcare Product Regulatory Agency (MHRA, United Kingdom) and Prof. Piet Maes (Rega Institute Leuven, Belgium) for providing the authentic SARS-CoV-2 viruses. Additionally, we thank the team at IGA Technology Services and Davide Scaglione for their contribution of authentic virus amplicon sequencing and data analysis services. We also thank Kath Hardcastle and the Comparative Medicine team at Moderna for the planning and execution of the in vivo studies.

## Author contributions

Y.C. designed and supervised the study. F.J., A.Z.W., L.M.W. and Y.C. wrote the manuscript with inputs from all authors. L.F., L.W. and X.W. solved and analyzed the Cryo-EM structures. Y.Y. and Youchun W. constructed pseudoviruses. P.W. and F.J. performed the rVSV escape screening experiments and data analysis. J.H. performed the authentic virus escape assays and data analysis. L.Y. performed the SPR experiments. P.W., L.Y., T.X., Yao W. and F. S. performed the pseudovirus neutralization assays. W.S., X.N., R.A. and Y.W. isolated the mAbs. J.W. (Changping Laboratory), L.L, L.Y. and F.S. performed protein expression and purification experiments. J.W. (BIOPIC) and F.J. analyzed the DMS data. S.Y., L.F., and F.J. performed sequence analysis and illustration. A.Z.W., S.P., and L.M.W. conceptualized and planned the mRNA delivery study. K.W., D.M.B., D.L., and T.S. performed the mouse serum neutralization assays and data analysis. C.H., L.M., and T.K. performed and supervised the pharmacokinetic ELISA assays.

## Extended Data Figures

**Extended Data Fig. 1.**
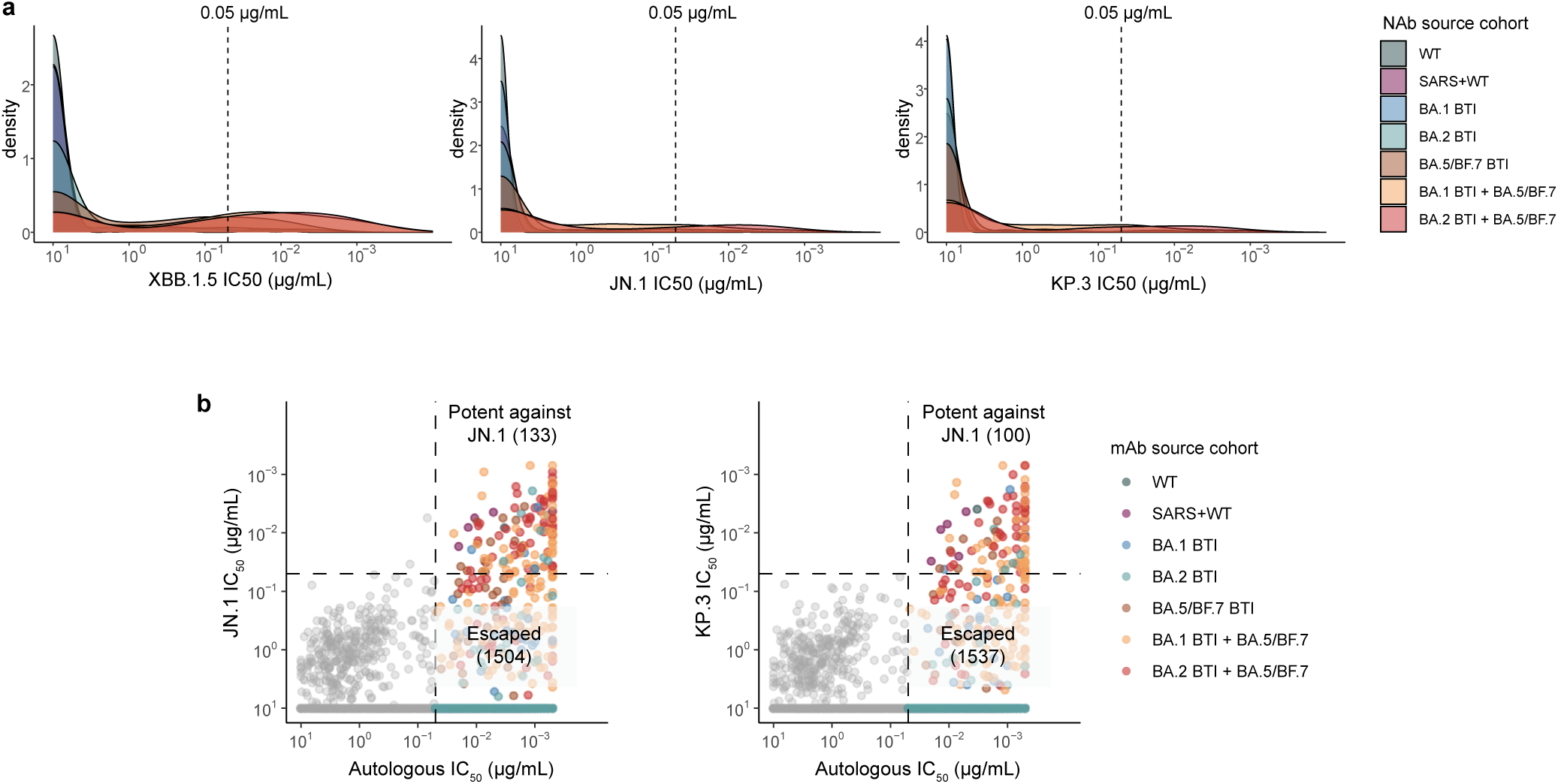
Neutralization distribution of mAbs collected. **a**, Distribution of the neutralization activities (IC_50_) of potent autologous NAbs against XBB.1.5 and JN.1. **b**, Scatter plots showing the relationship between the autologous neutralization activities and JN.1/KP.3-neutralizing activities of the isolated mAbs.

**Extended Data Fig. 2.**
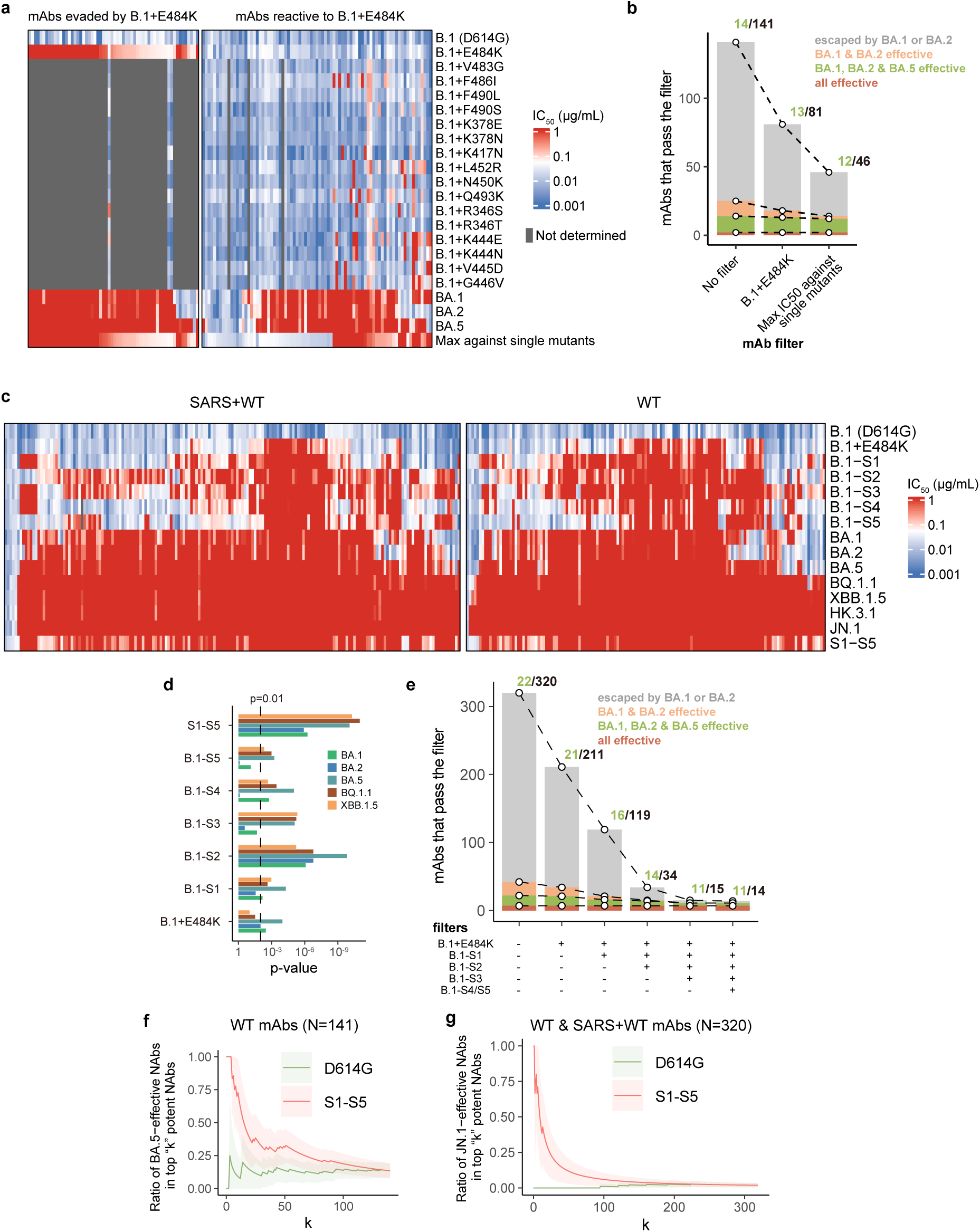
Neutralization activities for the identification of bnAbs. **a**, Heatmap of the neutralization activities (IC_50_) for the mAbs from SARS-CoV-2 WT cohort against the designed single mutants and Omicron variants. **b**, Number of NAbs from WT cohort that pass the filter of designed single mutants. Ratio of BA.1, BA.2, and BA.5-potent NAbs among the passed NAbs are annotated above the bar of each combination of filter. **c**, Heatmap of the neutralization activities (IC_50_) for the mAbs from early cohorts (SARS+WT and WT) against the designed mutants and real-world Omicron variants. “S1-S5” indicates the highest IC_50_ against the five designed mutants. **d**, Significance for the enrichment of BA.1, BA.2, BA.5, BQ.1.1, or XBB.1.5-potent NAbs within NAbs that were from WT or SARS+WT cohort pass each filter of designed mutants (two-sided hypergeometric test). **e**, Number of NAbs from WT or SARS+WT cohort that pass the filter of designed mutants. Ratio of BA.1, BA.2, and BA.5-potent NAbs among the passed NAbs are annotated. **f-g**, Ratio of BA.5 or JN.1-potent NAbs within the NAbs from WT cohort (f), or WT in addition to SARS+WT cohort (g) with “top k” neutralization activities against D614G or S1-S5. The error bars indicate 95% confidence interval under normal distribution.

**Extended Data Fig. 3.**
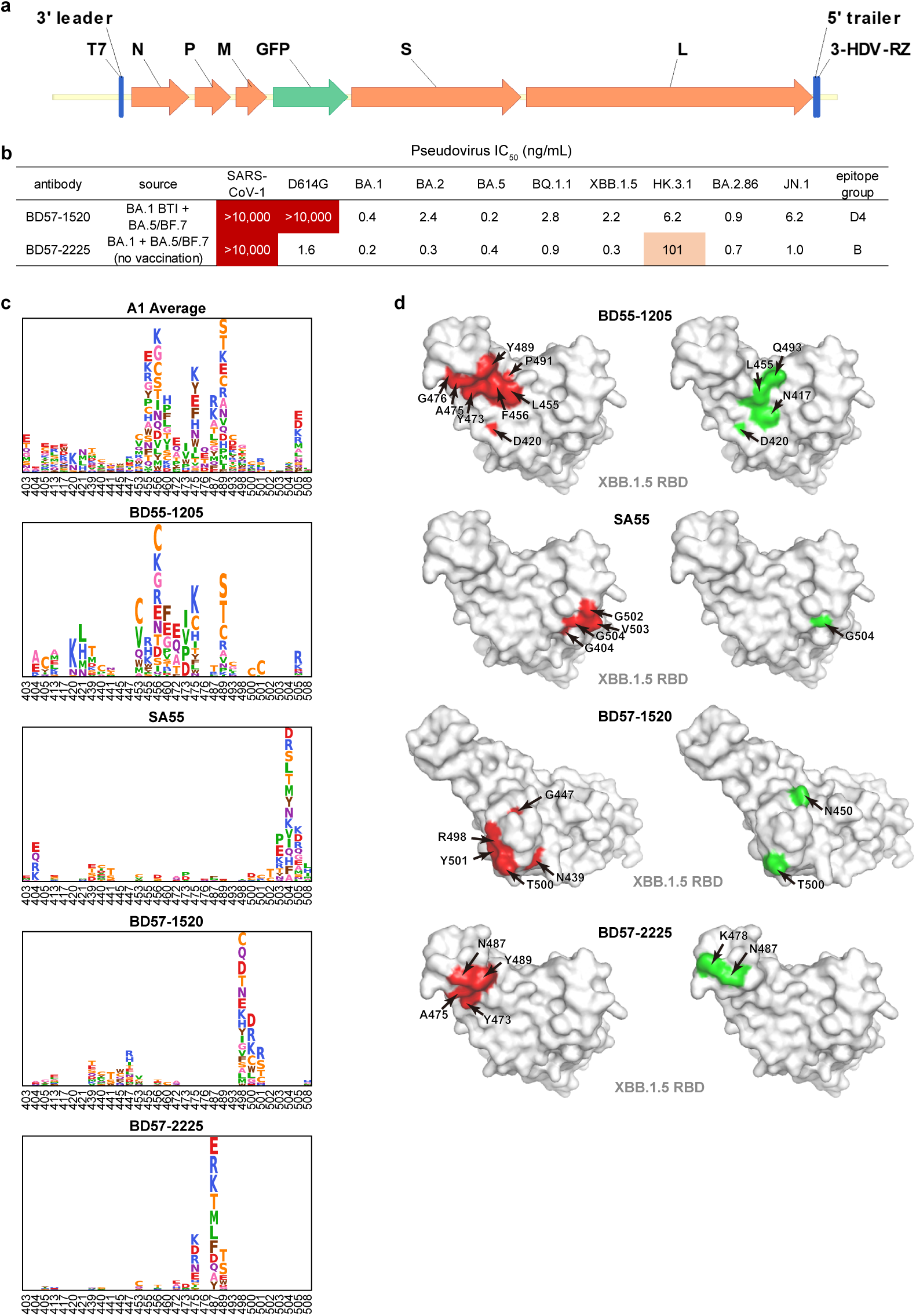
DMS and rVSV screening indicate the epitopes targeted by bnAbs. **a**, Schematic for the construction of the SARS-CoV-2 XBB.1.5 Spike-pseudotyped rVSV genome. **b**, Information of the two non-competing NAbs utilized in the rVSV screening assays. **c**, DMS escape profiles (based on BA.5 RBD) of the NAbs involved in the rVSV assays. The average profile of antibodies in epitope group A1 is also shown for comparison with BD55-1205. **d**, Key RBD sites that may be involved in the binding of NAbs (BD55-1205, SA55, BD57-1520 and BD57-2225) as determined by rVSV screening (green) and DMS (red) are marked on the structural model of XBB.1.5 RBD (PDB: 8WRL).

**Extended Data Fig. 4.**
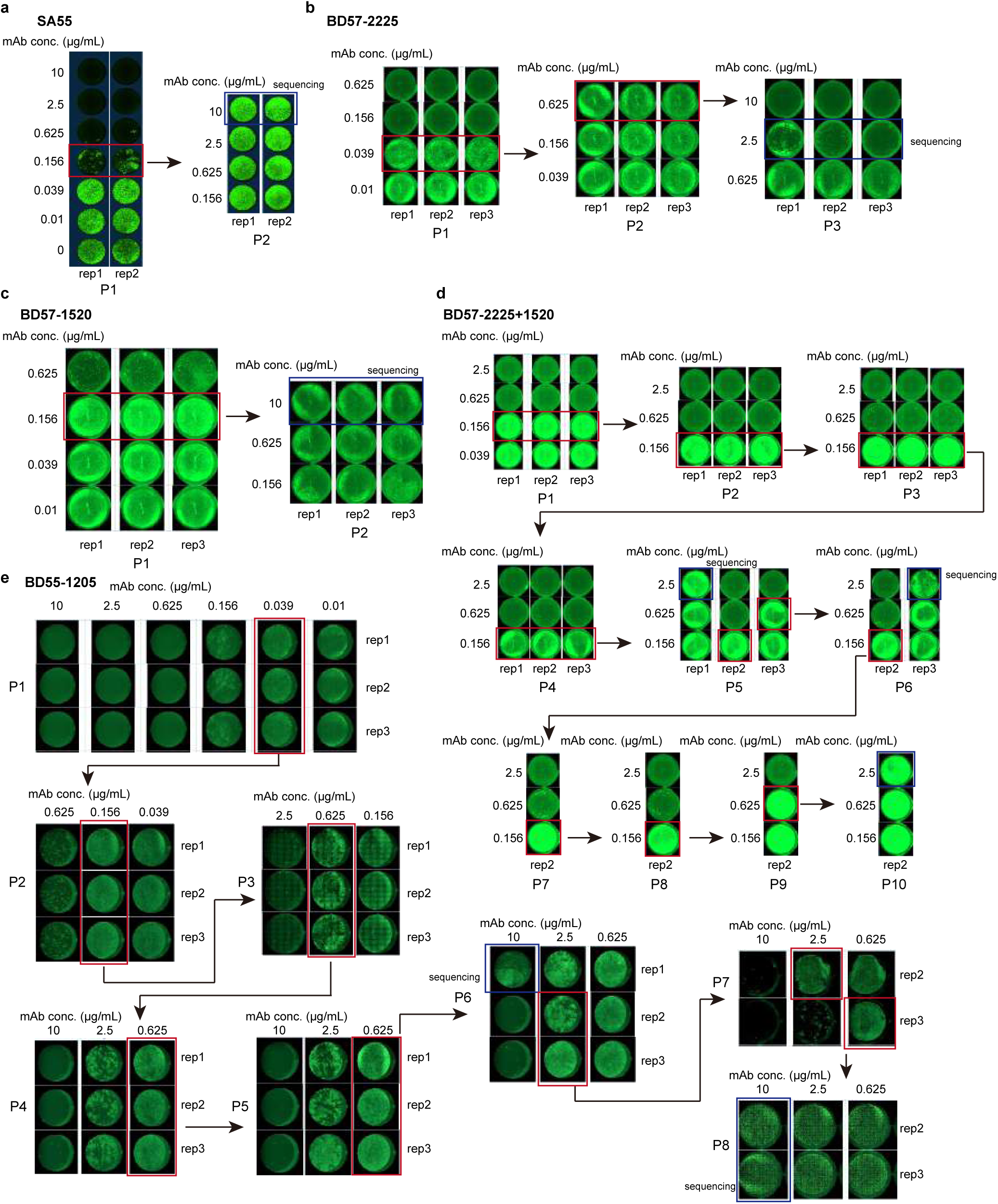
Raw images of the rVSV escape mutants screening assays. Raw images of the rVSV passages under the pressure of SA55 (a), BD57-2225 (b), BD57-1520 (c), BD57-2225+1520 (d), and BD55-1205 (e). Red rectangles indicate the well for the next passage, and blue rectangles indicate the well for Sanger sequencing. All of the experiments were performed on 24-well plates. The diameter of each well is 18 mm.

**Extended Data Fig. 5.**
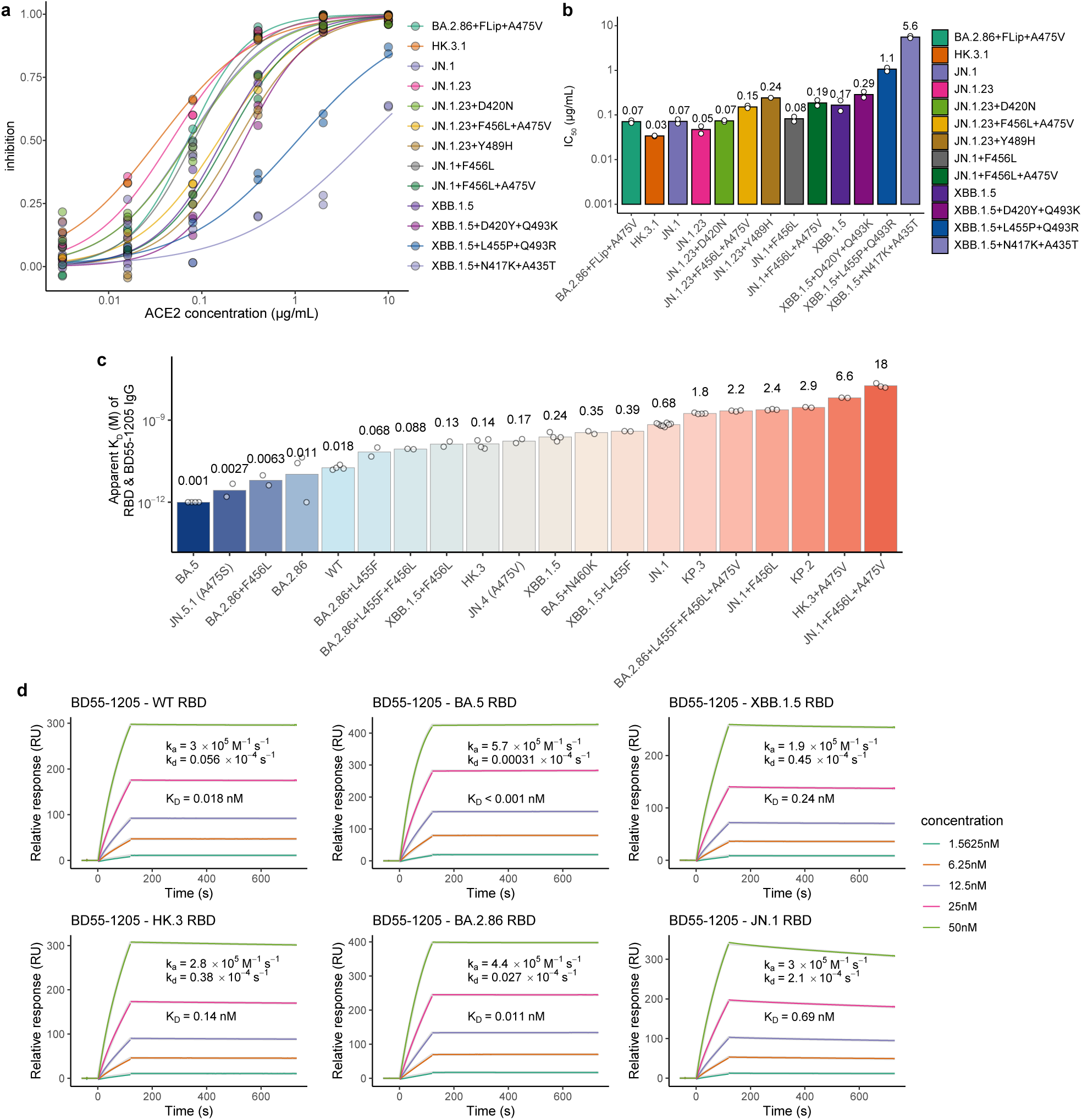
SARS-CoV-2 variant RBD-binding affinity of BD55-1205 and hACE2. **a**, Inhibition curves of soluble hACE2 against SARS-CoV-2 Omicron variant pseudoviruses. **b**, IC_50_ of soluble hACE2 against the variants. Geometric mean values are shown and annotated above the bars, and the circles indicate each of two biological replicates. **c**, RBD apparent binding affinity of BD55-1205 IgG to SARS-CoV-2 RBD variants determined by SPR assays. Geometric mean of apparent K_D_ (nM) is shown and annotated above the bars. Each circle indicates one independent experiment and the number of circles indicate the sample sizes. **d,** SPR sensorgrams of the binding of BD55-1205 to six major SARS-CoV-2 RBD variants. The association and dissociation kinetic coefficients (k_a_, k_d_), and the apparent dissociation equilibrium constant (K_D_) are annotated.

**Extended Data Fig. 6.**
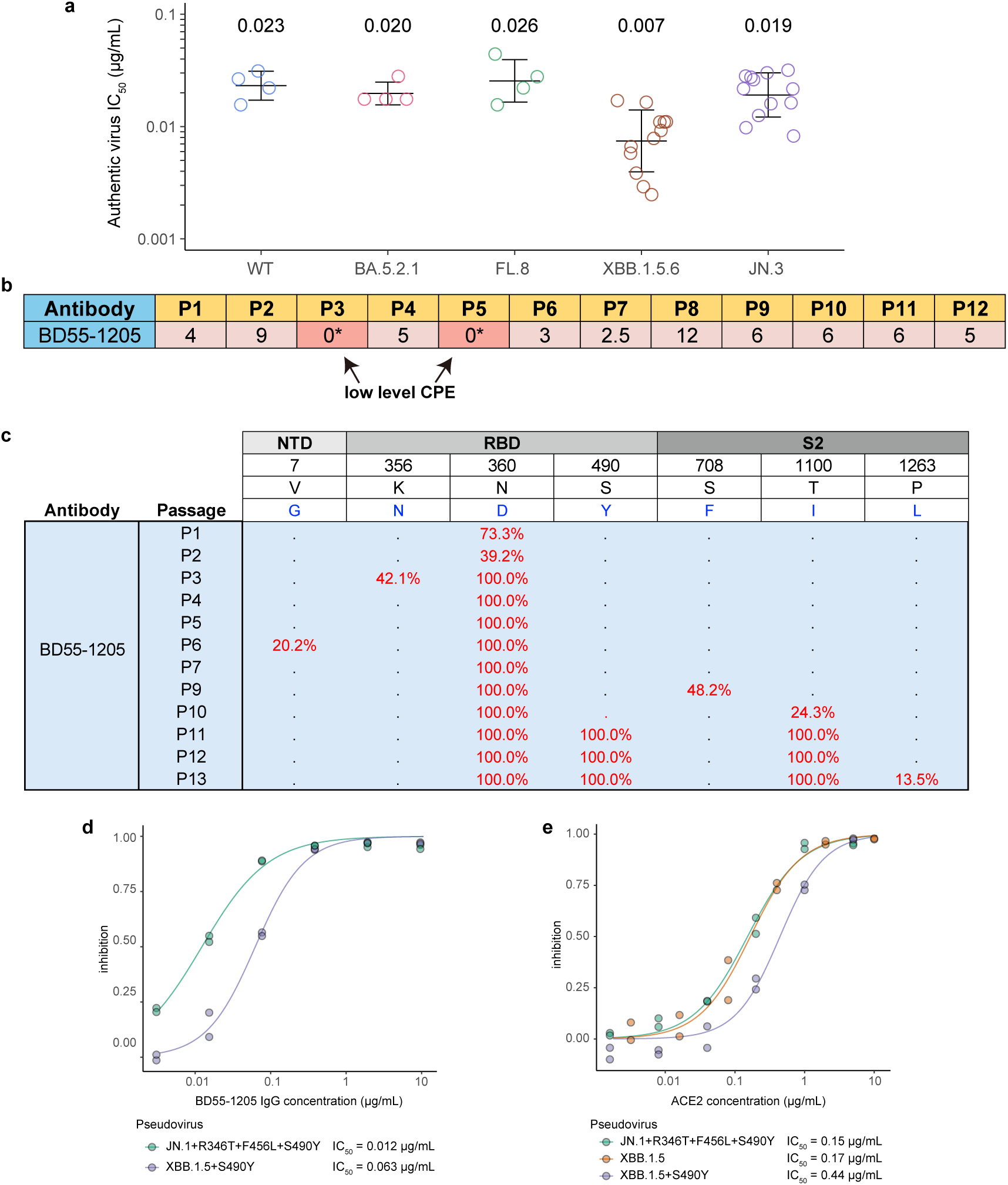
BD55-1205 inhibits SARS-CoV-2 authentic virus. **a**, IC_50_ of BD55-1205 against authentic SARS-CoV-2 isolates WT (n=4), BA.5.2.1 (n=4), FL.8 (n=4), XBB.1.5.6 (n=12), and JN.3 (n=12). Geometric mean IC_50_ (μg/mL) is labeled above the points and geometric SD is indicated by error bars. Each circle indicates a biological replicate. **b**, Number of wells in the plate that show protection from CPE in each passage in the escape screening assay using XBB.1.5.6 authentic virus. Asterisks indicate low-level/ambiguous CPE observation. **c**, Spike mutations observed in viral S protein resolved using amplicon-based deep sequencing of viral genome at each passage with BD55-1205 selection. **d-e**, Neutralization curves of BD55-1205 (**d**) or soluble hACE2 (**e**) against SARS-CoV-2 Omicron variant pseudoviruses with Spike S490Y mutation. XBB.1.5 inhibition curve is shown here again for comparison. The circles indicate results of two independent replicates.

**Extended Data Fig. 7.**
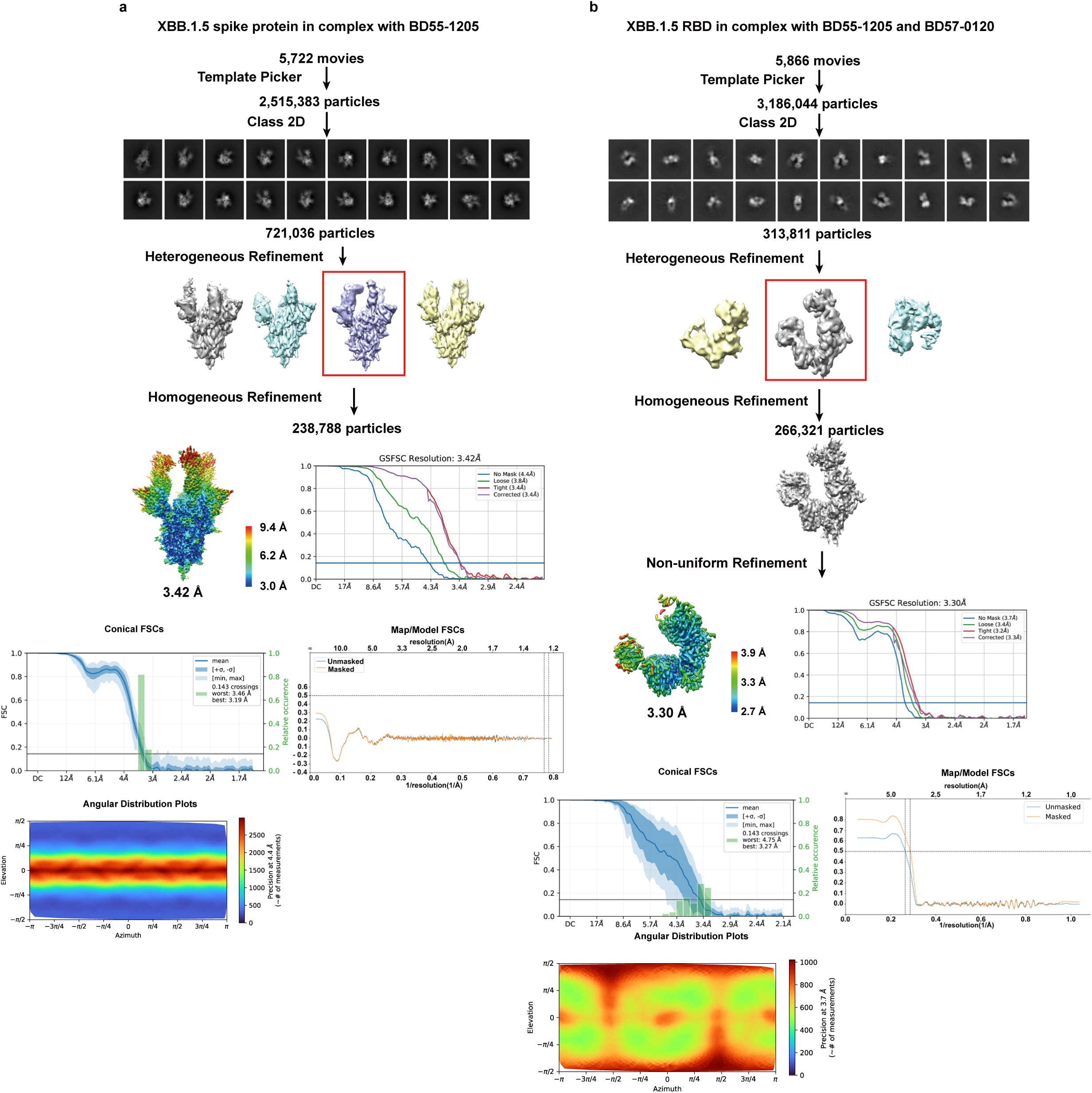
Workflow for the processing of Cryo-EM data. **a**, Workflow for the processing of raw Cryo-EM images for XBB.1.5 Spike in complex of BD55-1205. **b**, Workflow for the processing of raw Cryo-EM images for XBB.1.5 RBD in complex of BD55-1205 and BD57-0120.

**Extended Data Fig. 8.**
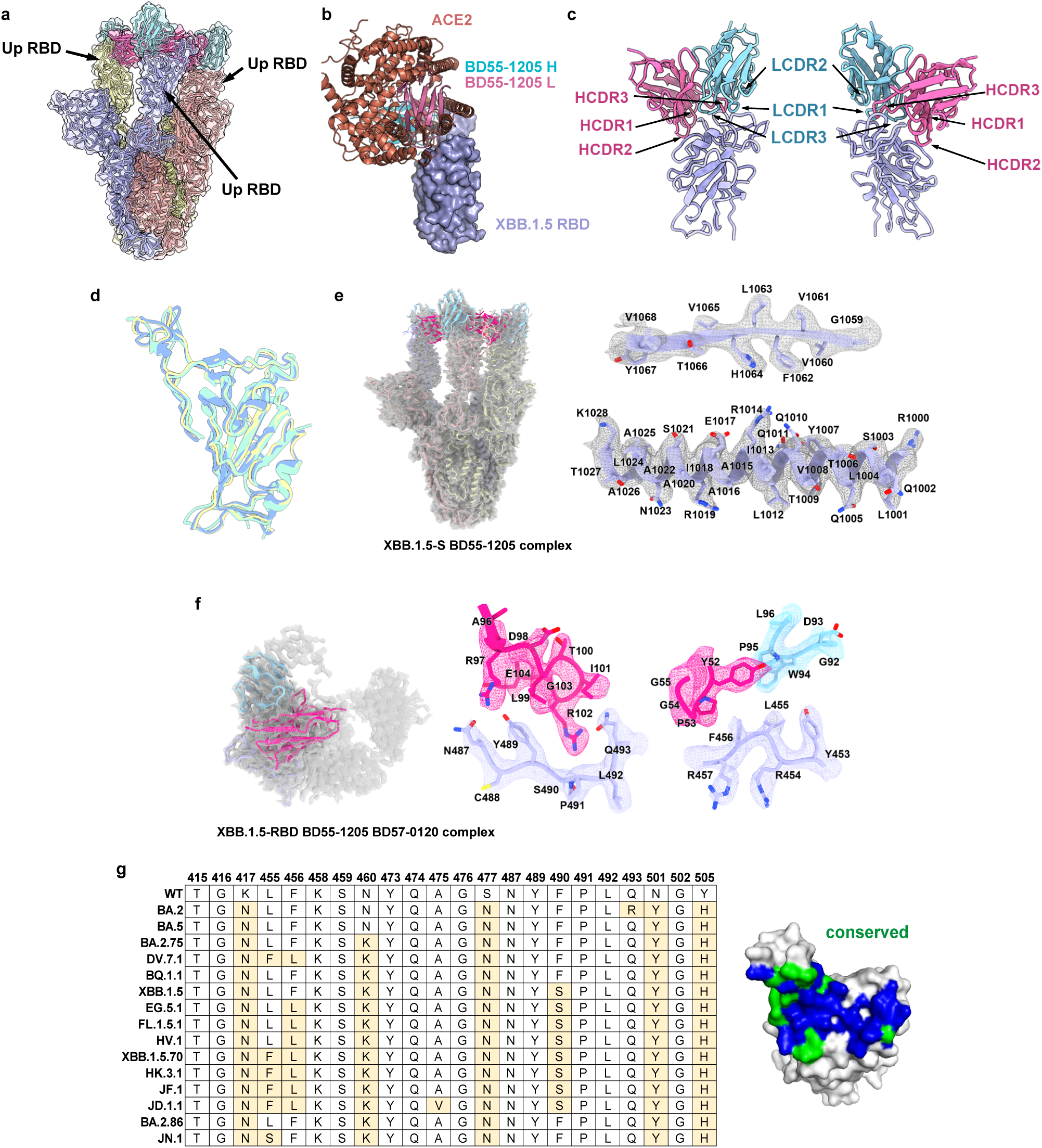
Structure of BD55-1205 in complex with Spike and RBD. **a**, Structure of BD55-1205 in complex of XBB.1.5 Spike with 3 RBD “up”. **b**, The epitope targeted by BD55-1205 is highly similar to hACE2 receptor binding sites. **c**, BD55-1205 interacts with XBB.1.5 RBD via heavy chain and light chain CDRs. **d**, Comparison of XBB.1.5 spike protein complex, RBD-BD55-1205-BD57-0120 complex cryo-EM structure, and RBD crystal structure. The RBD from XBB.1.5 spike protein complex (blue, this study), RBD crystal structure (green, PDB 9AU1) were superposed onto RBD double Fab cryo-EM structure (yellow, this study) respectively. The RMSD between RBD crystal structure and double Fab cryo-EM structure is 0.798 Å, and the RMSD between RBD from spike-Fab complex and spike double Fab cryo-EM structure is 0.845 Å. **e**, Cryo-EM density for XBB.1.5 S in complex with BD55-1205 complex and selected β-sheet regions (residues 1059-1067) and α helix (residues 1000-1028). **f**, Overall map of XBB.1.5 RBD in complex with BD55-1205 and BD57-0120 Fab, and representative interaction interfaces are shown. RBD, heavy chain, and light chain are colored in blue, magenta, and cyan, respectively. **g**, RBD residues in SARS-CoV-2 variants targeted by BD55-1205. Conserved residues are marked in green, and the other residues targeted by BD55-1205 are marked in blue.

**Extended Data Fig. 9.**
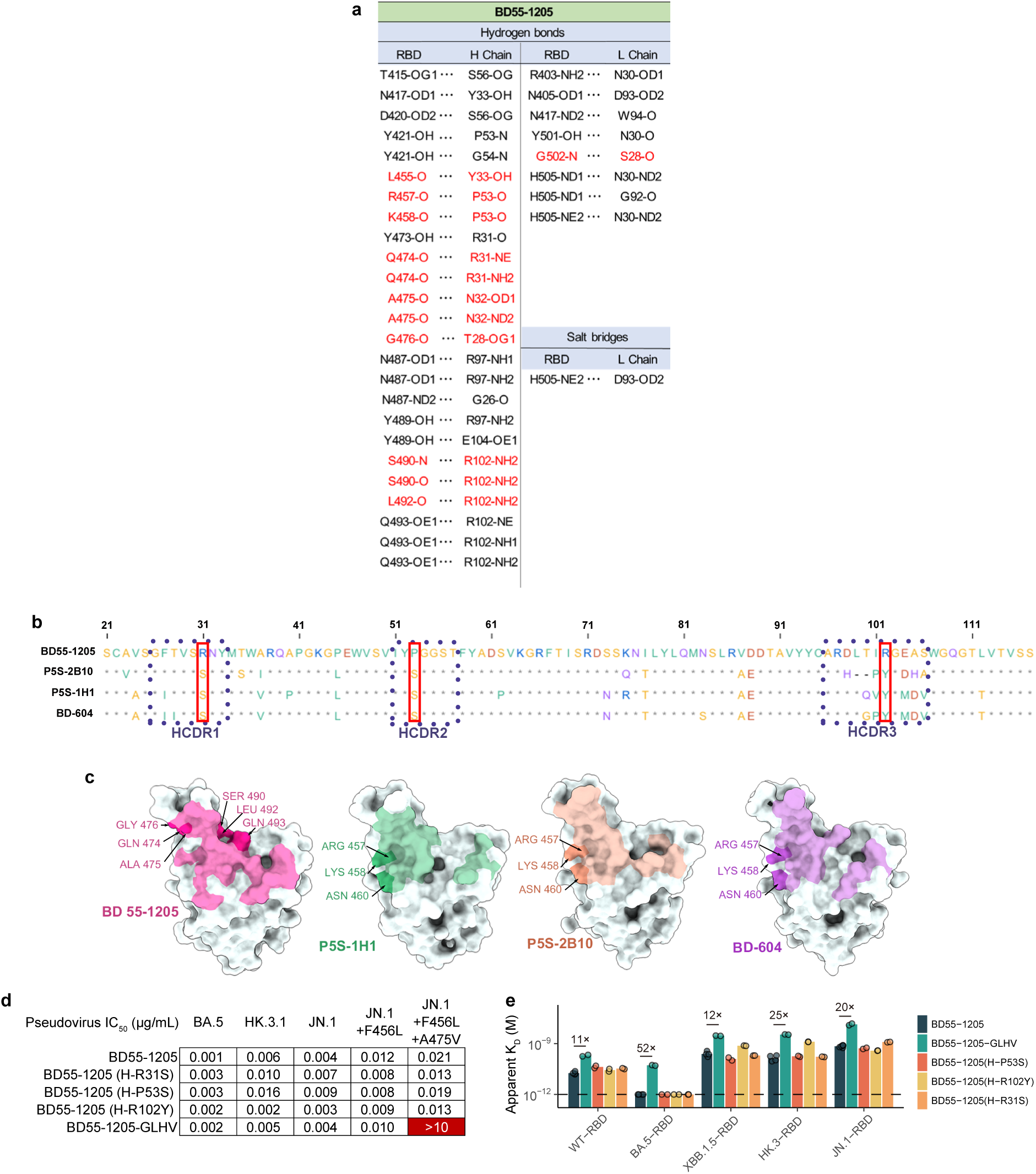
Comparison of BD55-1205 and Class 1 Nabs. **a**, A list of all polar interactions between SARS-CoV-2 variant RBD and BD55-1205. Interactions involving RBD backbone atoms are marked in red. **b**, Alignment of the heavy chains of BD55-1205, P5S-2B10, P5S-1H1, and BD-604. Heavy chain CDRs are marked in blue rectangles. Potential key mutations of BD55-1205 are marked in red rectangles. **c**, Footprints of the four antibodies on SARS-CoV-2 RBD. **d**, Neutralization of BD55-1205 with mutations on the heavy chain. **e**, RBD-binding affinity of BD55-1205 with mutations on the heavy chain. Geometric mean values are shown as bars, and each circle indicates one of two independent replicates.

**Extended Data Fig. 10.**
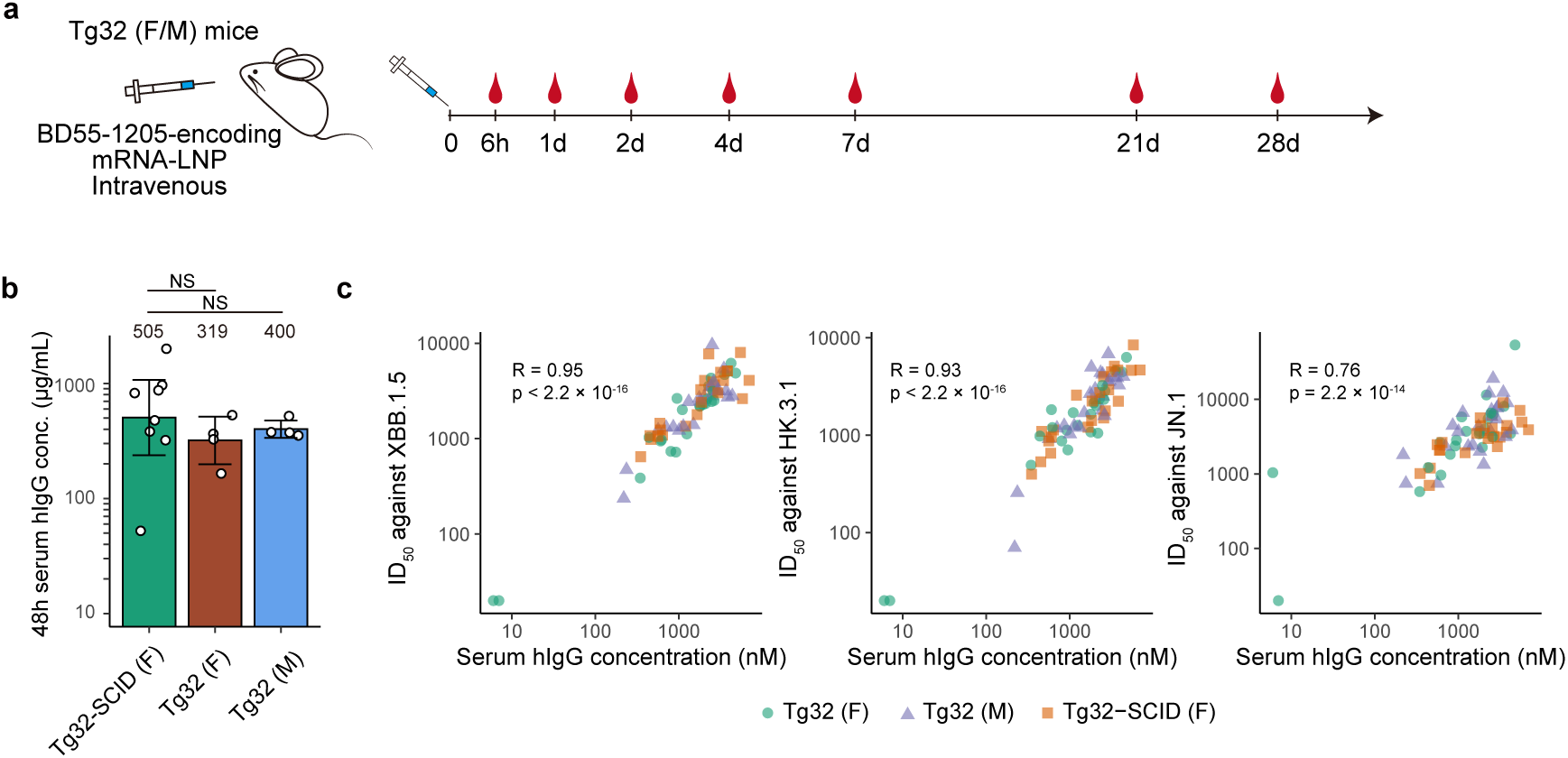
Immunization of Tg32 mice without SCID. **a**, A schematic of the experimental design for delivery of BD55-1205 in Tg32 mice without SCID. Female (F) or male (M) mice, 4 per group, received 0.5 mg/kg dose by intravenous injection on day 0 and serum was collected at the indicated time points. **b**, Peak serum concentration, occurring at 48 hours post LNP administration from the two SCID groups (n=8), and the non-SCID F (n=4) and M (n=4) group. Bar height and number above the bar indicate the geometric mean; error bars indicate 95% confidence intervals; empty symbols indicate individual animals. NS, not significant (two-sided Wilcoxon rank-sum test). **c**, Scatter plots showing the correlation between serum hIgG concentrations and the ID_50_ against the three variants at indicated timepoints. Pearson correlation coefficients (R) and the corresponding significance p-values are annotated (two-sided t-test).

## Supplementary Tables

**Table S1 | Information of SARS-CoV-2 RBD-specific mAbs involved in this study.** Immune history of source human donors, neutralization activities against major SARS-CoV-2 variants, and Fv sequences of SARS-CoV-2 RBD-targeting mAbs.

## Supplementary Information

Supplementary methods and statistics of Cryo-EM data

## References

1 Sachs, J. D. et al. The Lancet Commission on lessons for the future from the COVID-19 pandemic. Lancet 400, 1224–1280 (2022). 10.1016/S0140-6736(22)01585-9

2 Carabelli, A. M. et al. SARS-CoV-2 variant biology: immune escape, transmission and fitness. Nature Reviews Microbiology 21, 162–177 (2023). 10.1038/s41579-022-00841-7

3 Kaku, Y. et al. Antiviral efficacy of the SARS-CoV-2 XBB breakthrough infection sera against omicron subvariants including EG.5. The Lancet Infectious Diseases 23, e395–e396 (2023). 10.1016/S1473-3099(23)00553-4

4 Jian, F. et al. Convergent evolution of SARS-CoV-2 XBB lineages on receptor-binding domain 455–456 synergistically enhances antibody evasion and ACE2 binding. PLOS Pathogens 19, e1011868 (2023). 10.1371/journal.ppat.1011868

5 Kaku, Y. et al. Virological characteristics of the SARS-CoV-2 JN.1 variant. Lancet Infect Dis (2024). 10.1016/S1473-3099(23)00813-7

6 Yang, S. et al. Fast evolution of SARS-CoV-2 BA.2.86 to JN.1 under heavy immune pressure. Lancet Infect Dis (2023). 10.1016/S1473-3099(23)00744-2

7 Wang, Q. et al. XBB.1.5 monovalent mRNA vaccine booster elicits robust neutralizing antibodies against emerging SARS-CoV-2 variants. bioRxiv, 2023.2011.2026.568730 (2023). 10.1101/2023.11.26.568730

8 Dougan, M. et al. Bamlanivimab plus Etesevimab in Mild or Moderate Covid-19. N Engl J Med 385, 1382–1392 (2021). 10.1056/NEJMoa2102685

9 Gupta, A. et al. Early Treatment for Covid-19 with SARS-CoV-2 Neutralizing Antibody Sotrovimab. New England Journal of Medicine 385, 1941–1950 (2021). 10.1056/NEJMoa2107934

10 Weinreich, D. M. et al. REGN-COV2, a Neutralizing Antibody Cocktail, in Outpatients with Covid-19. New England Journal of Medicine 384, 238–251 (2020). 10.1056/NEJMoa2035002

11 Kelley, B. Developing therapeutic monoclonal antibodies at pandemic pace. Nature Biotechnology 38, 540–545 (2020). 10.1038/s41587-020-0512-5

12 Bruel, T. et al. Serum neutralization of SARS-CoV-2 Omicron sublineages BA.1 and BA.2 in patients receiving monoclonal antibodies. Nat Med 28, 1297–1302 (2022). 10.1038/s41591-022-01792-5

13 Cao, Y. et al. Omicron escapes the majority of existing SARS-CoV-2 neutralizing antibodies. Nature 602, 657–663 (2022). 10.1038/s41586-021-04385-3

14 Cao, Y. et al. BA.2.12.1, BA.4 and BA.5 escape antibodies elicited by Omicron infection. Nature 608, 593–602 (2022). 10.1038/s41586-022-04980-y

15 Iketani, S. et al. Antibody evasion properties of SARS-CoV-2 Omicron sublineages. Nature 604, 553–556 (2022). 10.1038/s41586-022-04594-4

16 Cox, M. et al. SARS-CoV-2 variant evasion of monoclonal antibodies based on in vitro studies. Nature Reviews Microbiology 21, 112–124 (2023). 10.1038/s41579-022-00809-7

17 Cameroni, E. et al. Broadly neutralizing antibodies overcome SARS-CoV-2 Omicron antigenic shift. Nature 602, 664–670 (2022). 10.1038/s41586-021-04386-2

18 Park, Y. J. et al. Antibody-mediated broad sarbecovirus neutralization through ACE2 molecular mimicry. Science 375, 449–454 (2022). 10.1126/science.abm8143

19 Luo, S. et al. An Antibody from Single Human VH-rearranging Mouse Neutralizes All SARS-CoV-2 Variants Through BA.5 by Inhibiting Membrane Fusion. Sci Immunol 0, eadd5446 (2022). 10.1126/sciimmunol.add5446

20 Nutalai, R. et al. Potent cross-reactive antibodies following Omicron breakthrough in vaccinees. Cell 185, 2116–2131 e2118 (2022). 10.1016/j.cell.2022.05.014

21 Hong, Q. et al. Molecular basis of receptor binding and antibody neutralization of Omicron. Nature 604, 546–552 (2022). 10.1038/s41586-022-04581-9

22 Westendorf, K. et al. LY-CoV1404 (bebtelovimab) potently neutralizes SARS-CoV-2 variants. Cell Rep 39, 110812 (2022). 10.1016/j.celrep.2022.110812

23 Bianchini, F. et al. Human neutralizing antibodies to cold linear epitopes and subdomain 1 of the SARS-CoV-2 spike glycoprotein. Science Immunology 8, eade0958 (2023). doi:10.1126/sciimmunol.ade0958

24 Liu, L. et al. Antibodies targeting a quaternary site on SARS-CoV-2 spike glycoprotein prevent viral receptor engagement by conformational locking. Immunity 56, 2442–2455.e2448 (2023). 10.1016/j.immuni.2023.09.003

25 Cao, Y. et al. Imprinted SARS-CoV-2 humoral immunity induces convergent Omicron RBD evolution. Nature 614, 521–529 (2023). 10.1038/s41586-022-05644-7

26 Li, G., Hilgenfeld, R., Whitley, R. & De Clercq, E. Therapeutic strategies for COVID-19: progress and lessons learned. Nature Reviews Drug Discovery 22, 449–475 (2023). 10.1038/s41573-023-00672-y

27 Yisimayi, A. et al. Repeated Omicron exposures override ancestral SARS-CoV-2 immune imprinting. Nature 625, 148–156 (2024). 10.1038/s41586-023-06753-7

28 Cao, Y. et al. Rational identification of potent and broad sarbecovirus-neutralizing antibody cocktails from SARS convalescents. Cell Rep 41, 111845 (2022). 10.1016/j.celrep.2022.111845

29 Cao, Y. et al. Potent Neutralizing Antibodies against SARS-CoV-2 Identified by High-Throughput Single-Cell Sequencing of Convalescent Patients’ B Cells. Cell 182, 73–84 e16 (2020). 10.1016/j.cell.2020.05.025

30 Cao, Y. et al. Humoral immune response to circulating SARS-CoV-2 variants elicited by inactivated and RBD-subunit vaccines. Cell Res 31, 732–741 (2021). 10.1038/s41422-021-00514-9

31 Starr, T. N. et al. Deep Mutational Scanning of SARS-CoV-2 Receptor Binding Domain Reveals Constraints on Folding and ACE2 Binding. Cell 182, 1295–1310 e1220 (2020). 10.1016/j.cell.2020.08.012

32 Ju, B. et al. Infection with wild-type SARS-CoV-2 elicits broadly neutralizing and protective antibodies against omicron subvariants. Nature Immunology 24, 690–699 (2023). 10.1038/s41590-023-01449-6

33 Chen, Y. et al. Broadly neutralizing antibodies to SARS-CoV-2 and other human coronaviruses. Nature Reviews Immunology 23, 189–199 (2023). 10.1038/s41577-022-00784-3

34 Li, H. et al. Establishment of replication-competent vesicular stomatitis virus-based recombinant viruses suitable for SARS-CoV-2 entry and neutralization assays. Emerg Microbes Infect 9, 2269–2277 (2020). 10.1080/22221751.2020.1830715

35 Schmidt, F. et al. High genetic barrier to SARS-CoV-2 polyclonal neutralizing antibody escape. Nature 600, 512–516 (2021). 10.1038/s41586-021-04005-0

36 Jian, F. et al. Evolving antibody response to SARS-CoV-2 antigenic shift from XBB to JN.1. bioRxiv, 2024.2004.2019.590276 (2024). 10.1101/2024.04.19.590276

37 Taylor, A. L. & Starr, T. N. Deep mutational scanning of SARS-CoV-2 Omicron BA.2.86 and epistatic emergence of the KP.3 variant. bioRxiv, 2024.2007.2023.604853 (2024). 10.1101/2024.07.23.604853

38 Barnes, C. O. et al. SARS-CoV-2 neutralizing antibody structures inform therapeutic strategies. Nature 588, 682–687 (2020). 10.1038/s41586-020-2852-1

39 Yuan, M. et al. Structural basis of a shared antibody response to SARS-CoV-2. Science 369, 1119–1123 (2020). doi:10.1126/science.abd2321

40 Zemlin, M. et al. Expressed Murine and Human CDR-H3 Intervals of Equal Length Exhibit Distinct Repertoires that Differ in their Amino Acid Composition and Predicted Range of Structures. Journal of Molecular Biology 334, 733–749 (2003). 10.1016/j.jmb.2003.10.007

41 Tuekprakhon, A. et al. Antibody escape of SARS-CoV-2 Omicron BA.4 and BA.5 from vaccine and BA.1 serum. Cell 185, 2422–2433 e2413 (2022). 10.1016/j.cell.2022.06.005

42 Du, S. et al. Structurally Resolved SARS-CoV-2 Antibody Shows High Efficacy in Severely Infected Hamsters and Provides a Potent Cocktail Pairing Strategy. Cell 183, 1013–1023 e1013 (2020). 10.1016/j.cell.2020.09.035

43 Levin, M. J. et al. Intramuscular AZD7442 (Tixagevimab–Cilgavimab) for Prevention of Covid-19. New England Journal of Medicine 386, 2188–2200 (2022). 10.1056/NEJMoa2116620

44 Schmidt, P. et al. Antibody-mediated protection against symptomatic COVID-19 can be achieved at low serum neutralizing titers. Science Translational Medicine 15, eadg2783 (2023). doi:10.1126/scitranslmed.adg2783

45 Roopenian, D. C., Christianson, G. J. & Sproule, T. J. in Mouse Models for Drug Discovery: Methods and Protocols (eds Gabriele Proetzel & Michael V. Wiles) 93–104 (Humana Press, 2010).

46 August, A. et al. A phase 1 trial of lipid-encapsulated mRNA encoding a monoclonal antibody with neutralizing activity against Chikungunya virus. Nature Medicine 27, 2224–2233 (2021). 10.1038/s41591-021-01573-6

47 Kose, N. et al. A lipid-encapsulated mRNA encoding a potently neutralizing human monoclonal antibody protects against chikungunya infection. Science Immunology 4, eaaw6647 (2019). doi:10.1126/sciimmunol.aaw6647

48 O’Brien, M. P. et al. Subcutaneous REGEN-COV Antibody Combination to Prevent Covid-19. New England Journal of Medicine 385, 1184–1195 (2021). 10.1056/NEJMoa2109682

49 Ison, M. G. et al. Prevention of COVID-19 Following a Single Intramuscular Administration of Adintrevimab: Results From a Phase 2/3 Randomized, Double-Blind, Placebo-Controlled Trial (EVADE). Open Forum Infectious Diseases 10 (2023). 10.1093/ofid/ofad314

50 Deal, C. E. et al. An mRNA-based platform for the delivery of pathogen-specific IgA into mucosal secretions. Cell Reports Medicine 4, 101253 (2023). 10.1016/j.xcrm.2023.101253

51 Nie, J. et al. Establishment and validation of a pseudovirus neutralization assay for SARS-CoV-2. Emerg Microbes Infect 9, 680–686 (2020). 10.1080/22221751.2020.1743767

52 Andreano, E. et al. SARS-CoV-2 escape from a highly neutralizing COVID-19 convalescent plasma. Proceedings of the National Academy of Sciences 118, e2103154118 (2021). doi:10.1073/pnas.2103154118

53 Reed, L. J. & Muench, H. A SIMPLE METHOD OF ESTIMATING FIFTY PER CENT ENDPOINTS12. American Journal of Epidemiology 27, 493–497 (1938). 10.1093/oxfordjournals.aje.a118408

54 Brai, A. et al. Exploring the Implication of DDX3X in DENV Infection: Discovery of the First-in-Class DDX3X Fluorescent Inhibitor. ACS Medicinal Chemistry Letters 11, 956–962 (2020). 10.1021/acsmedchemlett.9b00681

55 Wu, X. et al. Development of a Bioluminescent Imaging Mouse Model for SARS-CoV-2 Infection Based on a Pseudovirus System. Vaccines 11, 1133 (2023).

56 Lawson, N. D., Stillman, E. A., Whitt, M. A. & Rose, J. K. Recombinant vesicular stomatitis viruses from DNA. Proceedings of the National Academy of Sciences 92, 4477–4481 (1995). doi:10.1073/pnas.92.10.4477

57 Cui, Z. et al. Structural and functional characterizations of infectivity and immune evasion of SARS-CoV-2 Omicron. Cell 185, 860–871 e813 (2022). 10.1016/j.cell.2022.01.019

58 Nelson, J. et al. Impact of mRNA chemistry and manufacturing process on innate immune activation. Science Advances 6, eaaz6893 (2020). doi:10.1126/sciadv.aaz6893

59 Sabnis, S. et al. A Novel Amino Lipid Series for mRNA Delivery: Improved Endosomal Escape and Sustained Pharmacology and Safety in Non-human Primates. Molecular Therapy 26, 1509–1519 (2018). 10.1016/j.ymthe.2018.03.010

60 Narayanan, E. et al. Rational Design and In Vivo Characterization of mRNA-Encoded Broadly Neutralizing Antibody Combinations against HIV-1. Antibodies 11, 67 (2022).

61 Richner, J. M. et al. Vaccine Mediated Protection Against Zika Virus-Induced Congenital Disease. Cell 170, 273–283.e212 (2017). 10.1016/j.cell.2017.06.040

62 Richner, J. M. et al. Modified mRNA Vaccines Protect against Zika Virus Infection. Cell 169, 176 (2017). 10.1016/j.cell.2017.03.016

63 Whitt, M. A. Generation of VSV pseudotypes using recombinant ΔG-VSV for studies on virus entry, identification of entry inhibitors, and immune responses to vaccines. Journal of Virological Methods 169, 365–374 (2010). 10.1016/j.jviromet.2010.08.006

